# A cortico-subcortical loop for motor control via the pontine reticular formation

**DOI:** 10.1101/2023.08.09.552594

**Authors:** Emília Bősz, Viktor M. Plattner, László Biró, Kata Kóta, Marco A. Diana, László Acsády

**Author notes:** Corresponding author, Laboratory of Thalamus Research, Institute of Experimental Medicine; Budapest, Szigony u. 43. 1083. Hungary. These authors contributed equally to this work (Emília Bősz, Viktor M. Plattner, László Biró).

## Abstract

Movement and locomotion are controlled by large neuronal circuits like the cortex-basal ganglia (BG)-thalamus loop. Inhibitory output of the BG loop can directly control movement via specialized connections with the brainstem. Whether other parallel loops with similar logic exist is presently unclear. Here we demonstrate that glycine transporter 2-positive (GlyT2+) cells of the pontine reticular formation (PRF) receive cortical inputs and in turn innervate the thalamus. Thalamus-projecting GlyT2+ cells innervate subcortical regions distinct from BG targets. Cortical cells co-innervate PRF/GlyT2+ cells and thalamus as in the BG loops. Cortex exerts strong excitatory control on PRF/GlyT2+ cells and these neurons powerfully inhibit their thalamic targets. Activation of thalamus projecting PRF/GlyT2+ cells leads to contralateral turning. These results demonstrate that the PRF is part of a cortico-subcortical loop that regulates motor activity parallel to the BG circuits. The cortico-PRF-thalamus loop can synergistically control turning with the BG loops via distinct descending pathways.

**Graphical abstract:** 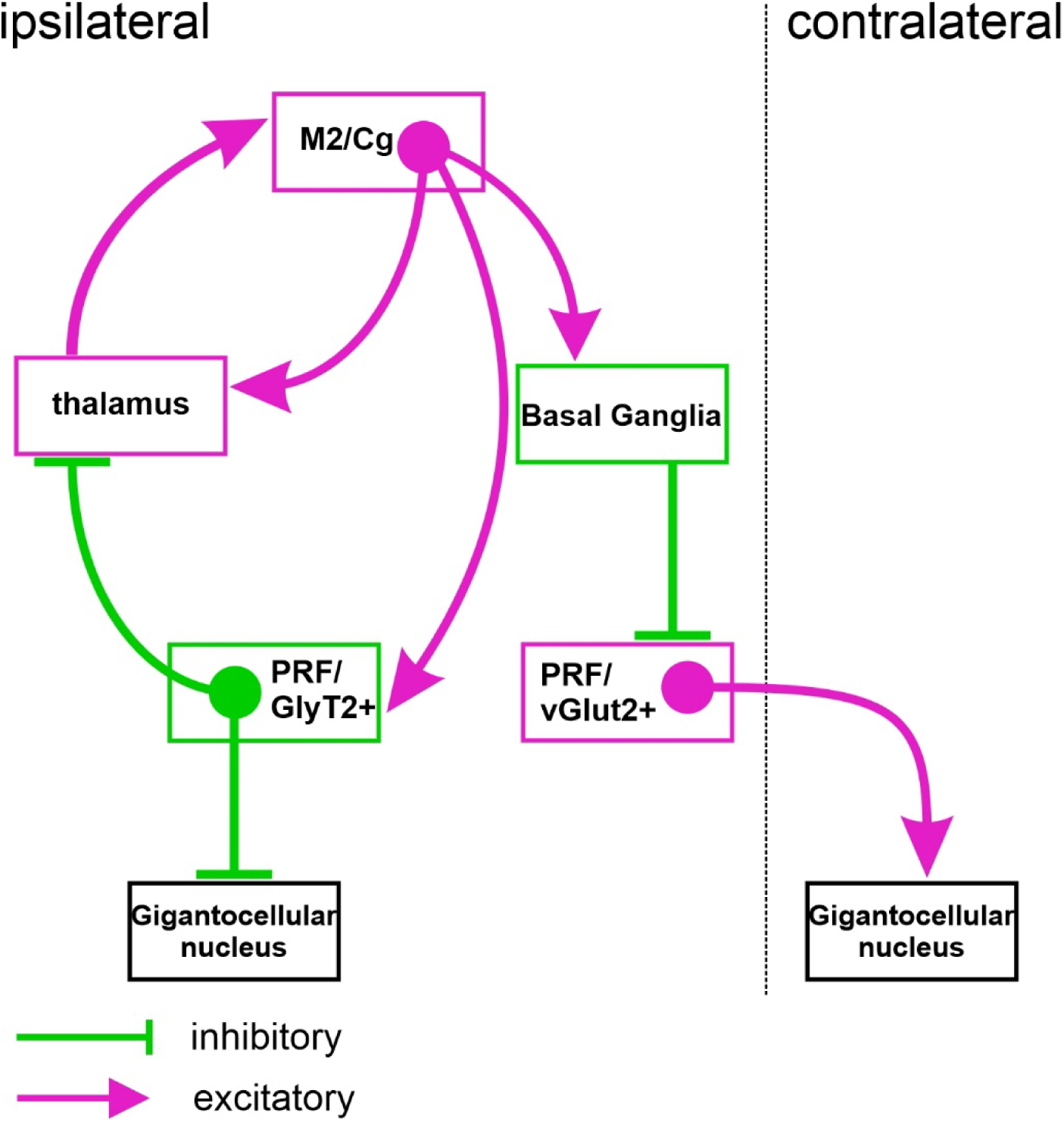

## Introduction

Voluntary movements are controlled via diverse cortical and subcortical centers^1,2^. Among these, the basal ganglia (BG) are one of the most intensively studied and are thought to be responsible for the planning, execution, and learning of complex motor sequences^2–5^. Two key features characterize the BG circuits. First, cortex, BG, and thalamus are engaged in the formation of complex loops ^6^–8; second, the output of these loops is communicated as a strong inhibitory signal to brainstem locomotor centers via the BG output nuclei ^9,10^.

The cortex-BG-thalamus loop can be regarded as a multiple-nested loop^11–13^ since the cortical signals return to the cortex via both the direct, thalamo-cortical projections and via an indirect loop that passes through the BG and the thalamus ^6–8^. Since the entire system is strongly topographical the indirect BG signals and the direct cortico-thalamic pathway are integrated in the same thalamic territory^6,8,14^. This pattern of connectivity is especially useful to generate multiple embedded rhythms, the hallmark of cortical and brainstem activity governing locomotion^1^. Individual cortical neurons can provide axon collaterals to both the direct thalamo-cortical pathway^15–17^ and the indirect cortex-BG loop ensuring that an identical cortical message is transferred via these two nested pathways.

How the cortex-BG-thalamus loop communicates with the brainstem locomotor centers has been poorly investigated until recently. By now it appears that the BG innervates multiple brainstem locomotor centers and can have a strong impact on their activity^9^. A recent study^10^ has demonstrated a specific role of the BG in regulating turning behavior via the pontine reticular formation (PRF, its rostral part also referred to as nucleus pontis oralis, PnO). PRF has already been implicated earlier in motor control^18,19^. Cregg et al.,^10^ showed that the substantia nigra pars reticulata (SNR), a major output nucleus of the BG, can effectively inhibit in the excitatory vesicular glutamate transporter 2 (vGLUT2)-containing neurons in the PRF. These PRF/vGLUT2+ neurons, in turn, innervate the contralateral gigantocellular nucleus (Gi) in the lower brainstem. Gi cells can mediate ipsiversive turning and motor arrest via their direct reticulo-spinal connections ^20,21^. The new study shows that decreased inhibitory SNR output results in increased PRF/vGLUT2+ output which consequently elevates contralateral Gi activity leading to a contralateral turning, relative to the side of SNR-PRF interaction. Increased BG output has the opposite effect^10^. Modulating the PRF/vGlut2+-Gi pathway restores turning ability after striatal damage, implying that impairment in this pathway could contribute to the disabling turning deficits associated with Parkinson’s disease^10^.These data demonstrate how the output of the cortex-thalamus-BG loop can have direct behavioral consequences via modulating the activity of the brainstem motor centers.

Curiously, the BG output avoided the other major cell type of the PRF, the equally numerous inhibitory, glycine transporter 2-expressing cells (PRF/GlyT2+ cells)^10,22^. At the same time, however, Gi neurons, ipsilateral to the turning, receive a strong inhibition^10^, the sources of which are unknown. PRF/GlyT2+ neurons have dual GABAergic/glycinergic phenotype and can innervate the same thalamic regions (the intralaminar/parafascicular nuclei, IL/Pf) as the BG-thalamic pathway^23^. IL/Pf have already been also implicated in movement coordination, turning and orienting behavior^24–26^. The thalamic projecting PRF/GlyT2+ cells establish large, highly effective multisynaptic inhibitory terminals, very similar to those of the BG terminals in the thalamus^23,27^. PRF is known to receive input from higher-order motor cortical areas^28,29^ and photoactivation of thalamus-projecting PRF/GlyT2+ cells results in motor arrest^23^. These data indicate that, even though PRF/GlyT2+ cells are not innervated directly by the BG output, they may participate in an independent cortico-thalamic loop and can contribute to motor control.

Based on these data, in this study, we tested whether thalamic projecting PRF/GlyT2+ neurons can represent a brainstem movement control system involved in a nested loop organization with the cortex and thalamus that can contribute to turning behavior. Our data demonstrate that thalamus-projecting PRF/GlyT2+ cells receive strong cortical excitatory inputs, via cortical L5 neurons. L5 neurons, in addition to innervating the PRF, also arborize in the IL/Pf, the target of PRF/GlyT2+cells. We also show that thalamic projecting PRF/GlyT2+ cells can evoke pronounced inhibitory action on their thalamic targets and affect turning behavior synergistically with the PRF/vGLUT2+ cells via a selective ipsilateral projection to Gi.

## Results

### M2/cingulate cortical neurons innervate PRF/GlyT2+ neurons

In order to visualize cortical inputs to PRF/GlyT2+ neurons, we used anterograde virus-mediated tracing in RBP4-Cre//GlyT2-eGFP double transgenic mice. RBP4 is known to be expressed in deep L5 pyramidal cells the project to subcortical targets^30^. We injected AAV5-DIO-ChR2-mCherry virus into the secondary motor/cingulate (M2/Cg) cortical area since these regions is known to contain the highest concentration of PRF projecting cells^23^ (Figure 1A-C). GlyT2+ cells were detected by the eGFP expression^22^. Anterogradely labeled M2/Cg fibers formed a profuse axon arbor within the entire rostral PRF (Figure 1D-F). The distribution of cortical afferents displayed significant overlap with that of the PRF/GlyT2+ neurons (Figure 1D-F) but was not homogeneous within the PRF. A density map of cortical afferents showed a dorsomedial to ventrolateral decrease in fiber density within the PRF/GlyT2+ zone (Figure 1G, S1A-C). This gradient was present at all coronal levels examined on averaged maximum intensity projection of the fiber density maps in 3 animals (Figures 1H1-3, S1A-C). Caudal injections into the M2/Cg cortex (n=3) resulted in weaker PRF innervation (Figure 1I). The quantitative analysis of intensity levels showed that the whole PRF was innervated by M2/Cg (Figure 1J). The posterior M2/Cg afferents displayed low-intensity levels (2-4) compared to the anterior M2/Cg innervation where we observed higher fiber density (intensity levels 1-7, Figure 1K).

**Figure 1:**
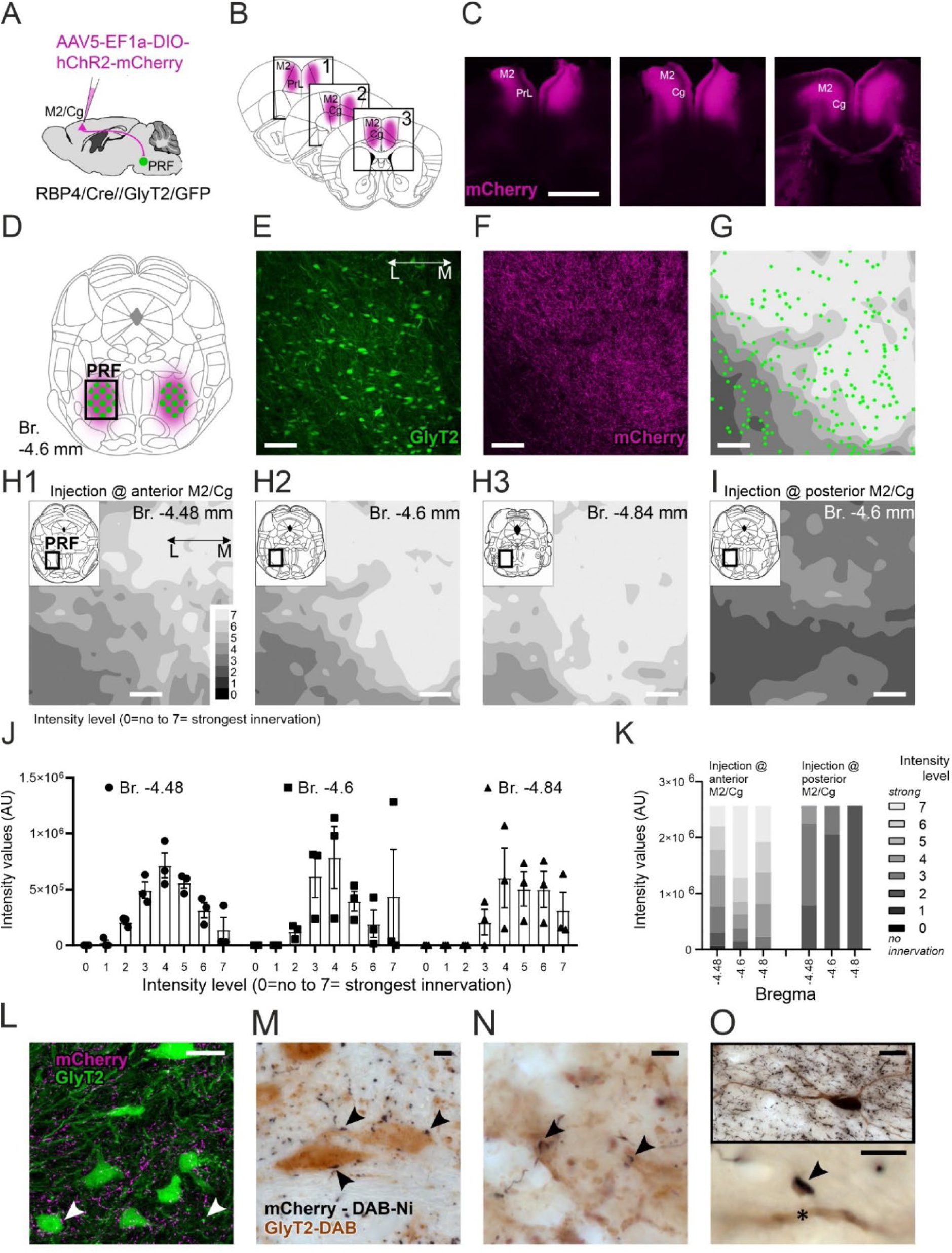
Cortical inputs to PRF/GlyT2+ cells. A) Scheme of anterograde tracing from the frontal cortical motor-related areas (M2/Cg) in the RBP4-Cre//GlyT2-eGFP mouse line (n=9 mice). B) Schematic view of the injection sites. C) Low-power fluorescent micrographs of the cortical injection sites at three anteroposterior levels. D) Schematic view of the M2/Cg fibers (magenta shade) and PRF/GlyT2+ cells (green dots). The black rectangle, the position of the micrographs and heatmaps in e-g. E-F) Confocal micrographs (n=5 mice) of the glycinergic cells (E) and anterogradely labeled M2/Cg cortical fibers (F) in the PRF glycinergic zone. G) Fiber density heat map showing the distribution of cortical fibers (grey shading) and PRF/GlyT2+ cells (green dots). Higher fiber density is indicated with light grey colors. H1-3) Maximum intensity Z projections of average cortical fiber heat maps (n=3 animals) showing anterior M2/Cg inputs at three anteroposterior PRF levels extending from Br. −4.6 to Br. −4.84 (from the Paxinos atlas; 250µm) I) Maximum intensity Z projections of cortical fiber heat maps (n=3 animals) showing posterior M2/Cg inputs. J) Quantitative analysis of the anterior M2/Cg fibers in the PRF at three coronal levels. Intensity levels represent fiber density values between 0 and 7. Where 0 indicates no innervation and 7 presents the strongest innervation. K) Proportion of innervation densities in the PRF after anterior (left) and posterior (right) M2/Cg injections at three coronal levels. L) High power confocal fluorescent image of anterogradely labeled M2/Cg fibers (magenta) and PRF/GlyT2+ neurons (60x, n=5 mice) M-O) High power light microscopic image (n=9 mice) of close apposition between M2/Cg fibers (black) and the somata (k), dendrites (l), or a spine (m) of PRF/GlyT2+ neurons. Inset in m displays the juxtacellularly labeled PRF/GlyT2+ cell; arrowheads, cortical inputs; asterisk, and spine. Scale bars: c) 1 mm; e-i) 100 μm; j) 20 μm; k-l) 5 μm; m) 5μm, 20μm in the inset.

Next, we visualized anterior M2/Cg fibers and PRF/GlyT2+ neurons using high-power confocal imaging and high-power light microscopy using DAB/Ni (black) and DAB (brown) chromogens. With both methods, cortical afferents were in close proximity to PRF/GlyT2+ elements (Figure 1L). We found putative synaptic contacts on the dendrites and occasionally on the somata of GlyT2+ neurons (Figure 1M, N). In one case the spine of a PRF/GlyT2+ neuron, filled using the juxtacellular labeling method (see below), was found to be contacted by an M2/Cg cortical terminal (Figure 1O).

In order to identify the synaptic contacts formed by M2/Cg terminals in the PRF, we analyzed the ultrastructure of the M2/Cg synapses at the electron microscopic level (Figure 2A-D). M2/Cg terminals were small to medium-sized (mean=0.47 µm^2^ SD=0.31, Figure 2B-D, S1D), contained 1-3 mitochondria, and established asymmetrical synapses with 1-2 active zones mainly with dendrites (range: 0.197-1.437 µm, mean=0.62 µm, SD=0.31, Figure 2B-D). The postsynaptic elements have variable diameters. Using double immunocytochemistry we identified synaptic contacts between virus-labeled cortical afferents and PRF/GlyT2+ neurons in 10 cases (Figure 2D1-4). In 9 cases the postsynaptic structure was mid-caliber dendrite in 1 case it was a PRF/GlyT2+ spine. These data show that the anterior M2/Cg provides a dense innervation to the PRF, which contains GlyT2+ neurons and forms conventional synaptic contacts mainly with the dendrites of GlyT2+ neurons in the PRF.

**Figure 2.**
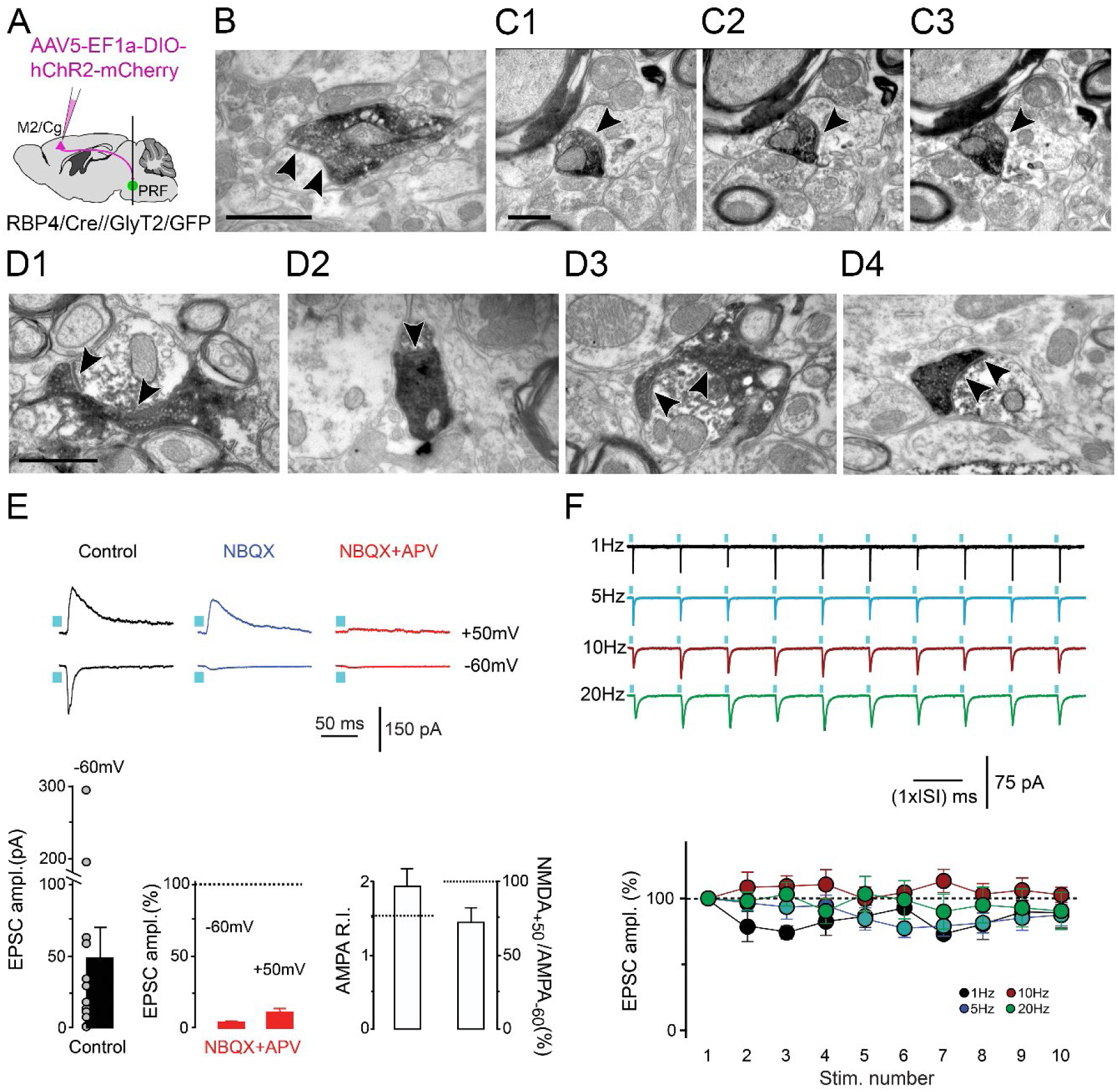
Cortical afferents target PRF/GlyT2+ cells and evoke a glutamatergic synaptic response. A) Scheme of anterograde tracing from the M2/Cg in the RBP4-Cre//GlyT2-eGFP mouse line. B-C) Electron micrographs of M2/Cg terminals (n=44, dark precipitates) in the PRF contacting mid-caliber dendrites. C1-C3) Serial EM images of the same axon terminal. Black arrowheads, synapses. D) Electron micrographs of M2/Cg terminals (DAB-Ni, dark, dense precipitate) establishing synapses on PRF/GlyT2+ dendrites (n=9, D1, D3, D4, DAB, light precipitate) and spines (n=1, D2). E) Optogenetically evoked (blue squares) AMPA and NMDA receptor-mediated components of the EPSCs elicited by M2/Cg terminals in the PRF. Traces from one exemplary experiment are shown above. Top: AMPA and NMDA components (n=16), control at −60 mV (n = 12; p = 0.002; Wilcoxon Signed-Rank Test) and +50 mV (n = 12; p = 0.003; Wilcoxon Signed-Rank Test). Below, the graphs represent the results from all the experiments. F) Top: Light-evoked EPSC-s displayed no significant depression during stimulation trains, at all tested frequencies ranging from 5 to 20 Hz. Bottom: 89.1 ± 11.5 % (n = 6; p > 0.05; Wilcoxon Signed-Rank Test), 87.3 ± 8.2 % (n = 14; p = 0.03); Wilcoxon Signed-Rank Test), 102.7 ± 5.7 % (n = 10; p = 0.7; Wilcoxon Signed-Rank Test), 90.5 ± 14.0 % (n = 12; p = 0.1; Wilcoxon Signed-Rank Test at 1 Hz, 5 Hz, 10 Hz and 20 Hz respectively. Source data are provided as a Source Data file. Data are represented as mean +/− SE. Scale bars: b) 1000 nm; c) 500 nm; d) 1000 nm.

### Synaptic properties of the cortical inputs in PRF/GlyT2+ neurons

Next, we investigated the physiology of synapses formed by the cortico-PRF terminals in GlyT2+ cells. In vitro, PRF slice preparations were made from RBP4-Cre//GlyT2-eGFP animals that received identical AAV5-DIO-ChR2-mCherry M2/Cg injections as in the case of the anterograde tracing experiments. Optogenetic activation with brief flashes (2-3 ms) of blue light evoked synaptic. At a recording potential of −60 mV, these triggered rapidly rising and decaying inward synaptic currents with an averaged amplitude amounting to 48.6 ± 20.2 pA; (Figure 2E, bottom panel). The amplitude of these responses was not modified by bath application of the GABAergic/glycinergic synaptic blockers SR95531 (2 µM) and strychnine (10 µM), thus demonstrating their excitatory nature. When recorded at a holding potential of +50 mV (see methods), the EPSCs showed clearly slower decaying kinetics than at −60 mV consistently with the appearance of an NMDA receptor-mediated component at the depolarized potential (Figure 2E, top panel, blue traces). Bath co-application of NBQX (10 µM) and DL-APV (50 µM), led to almost complete inhibition of the synaptic responses to 5.6 ± 1.4 % of the control at −60 mV, and to 10.1 ± 3.0 % at +50 mV (Figure 2E, top red traces), thus demonstrating the glutamatergic nature of the cortico-PRF/GlyT2+ inputs. The amplitude of the NMDA component, as measured at +50 mV, amounted to 68.7 ± 15.3 % of the AMPA component at −60 mV (Figure 2E). Moreover, the AMPA rectification index (measured as the ratio between the AMPA component amplitudes at −60 mV and at +50 mV) amounted to 1.9 ± 0.3 (n = 17; Figure 2E bottom), not distant from the theoretical value of 1.75 for a linear I-V curve (see Otsu et al., 2018). This strongly suggests that the AMPA receptors present at these synapses are calcium-impermeable in PRF/GlyT2+ neurons.

In control pharmacological conditions, we examined the dynamic behavior of the cortico-PRF glutamatergic responses with trains of 10 optogenetic stimulations at different frequencies (1 Hz, 5 Hz, 10 Hz and 20 Hz). Along the stimulation trains the synaptic responses in the PRF/GlyT2+ cells did not show a significantly depressing behavior, the amplitude of the responses to the 10^th^ stimulation being significantly smaller than the one to the first light pulse only at 5 Hz (see top traces in Figure 2F). With respect to the first optogenetic stimulation, the amplitudes of the responses to the 10^th^ light pulse were indeed 89.1 ± 11.5 %, 87.3 ± 8.2 %, 102.7 ± 5.7 %, 90.5 ± 14.0 % at 1 Hz, 5 Hz, 10 Hz, and 20 Hz, respectively (see bottom graph in Figure 2F). These data demonstrate that the cortical inputs to the PRF/GlyT2+ neurons identified morphologically exert a pronounced, conventional, purely glutamatergic synaptic action with little short-term depression.

### The impact of cortical activation on PRF/GlyT2+ neuronal activity

In order to assess the impact of cortical inputs on the spiking activity of PRF/GlyT2+ neurons *in vivo,* we recorded the evoked responses of the PRF cells in ketamine-xylazine anesthetized animals (n=10 animals, Figure 3). In 7 cases we injected the AAV5-DIO-ChR2-mCherry virus into M2/Cg in the RBP4-Cre//GlyT2-eGFP double transgenic mice and photostimulated the L5 neurons (Figure 3A-B). In 3 cases we applied bipolar electrical stimulation of the cortex in GlyT2/GFP transgenic mice (5 ms pulse width, 1-2 mA). In both stimulating conditions, we used trains of 1, 10, and 20 Hz stimulations (n=50 stimulus in 5 trains, Figure S2). To unequivocally identify and localize PRF/GlyT2+ neurons we used the juxtacellular recording and labeling methods^31^. Nine out of 10 neurons were recovered post hoc within the PRF and their GlyT2+ identity was determined by double fluorescent imaging (Figures 3C, and S2A).

**Figure 3.**
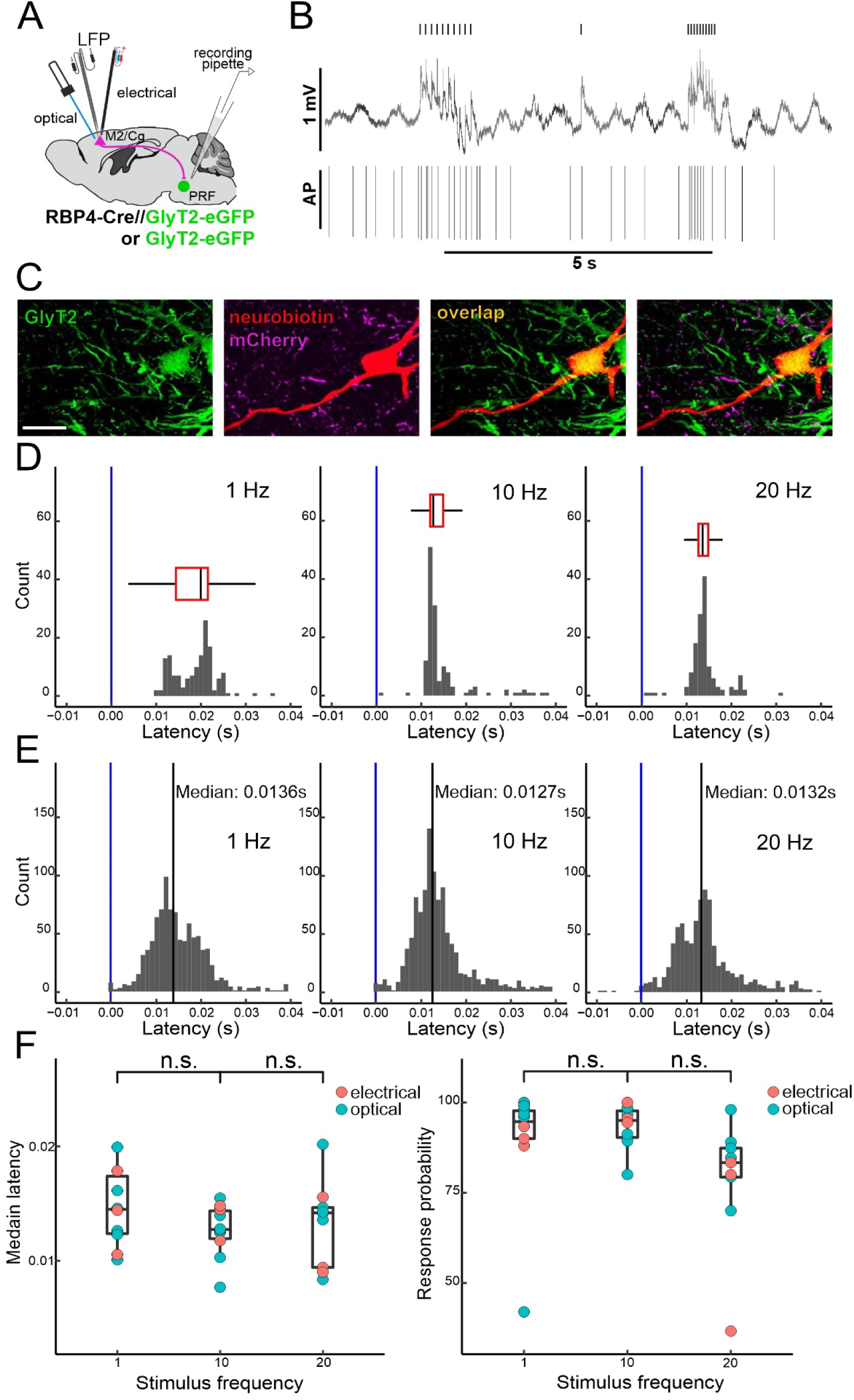
The effect of evoked cortical activity on PRF/GlyT2+ neurons. A) Scheme of the experiments (n=10 mice: 7 mice with optogenetic 5-ms-long pulses at 1-20 mW (RBP4-Cre//GlyT2-eGFP), 3 mice with electrical 5 ms pulses at 1-2 mA (GlyT2-eGPF)). B) Representative stimulus-evoked cortical field activity and evoked firing of a glycinergic PRF neuron. Small ticks on the top indicate laser stimulation. C) Representative fluorescent micrograph of a recorded and labeled PRF/GlyT2+ neuron. Green: GlyT2-eGFP; red: neurobiotin; magenta: M2/Cg fibers. D) Peristimulus time histograms (PSTH) of cortical stimulus-evoked firing in a representative PRF/GlyT2 neuron at 1, 10, and 20 Hz stimulation frequencies. Response probability and median latency are visualized by box plots.1 Hz: probability 92.86%, 12.5 ms peak +/− SEM; 10 Hz: probability: 92.86%, 12.6 ms peak +/− SEM, 20 Hz: probability 84.06%, 14.5 ms peak +/−SEM. E) Population PSTHs of juxtacellularly recorded and labeled PRF/GlyT2+ neurons (n=10) at 1, 10, and 20 Hz. 1 Hz median: 0.0136 s,10 Hz median: 0.0127 s, 20 Hz median: 0.0132 s. F) Median latency (left) 1 vs. 10 Hz, n.s. p=0.22; 10 Hz vs. 20 Hz n.s. p=0.3, Mood’s median test and response probability (right) 1 vs. 10 Hz, n.s p=0.59; 10 Hz vs. 20 Hz n.s. p=0.65 Mood’s median test of the PRF/GlyT2+ neurons. * 0.05<p; ** 0.01<p; *** p<0.001; n.s. - no significant difference. Source data are provided as a Source Data file Scale bars: c) 20 μm.

Cortical stimulation at 1 Hz generated reliable spike responses with short latency peaks, in all PRF/GlyT2+ cells (Figure 3D-E) using both stimulation methods. Increasing the stimulation frequency to 10 and 20 Hz resulted in similar high-fidelity, fast responses (Figure 3D-E). We found no significant difference between stimulation frequencies in response probability and latency (Figure 3F) indicating reliable signal transmission even at high presynaptic activity. To assess the alteration of firing output of the PRF/GlyT2+ cells within the stimulation train at various input frequencies, we calculated the normalized evoked AP within a 50 ms time window after every stimulus (Figure S2B). The data demonstrated that the spiking activity of individual PRF/GlyT2+ cells did not depress within the stimulus train irrespective of the stimulation frequencies (Figure S2B). These in vivo data confirm the in vitro physiological data and show that frontal cortical cells can evoke fast, reliable spike responses in PRF/GlyT2+ cells even at high presynaptic stimulation frequencies.

### Impact of spontaneous cortical activity on PRF/GlyT2+ cells

Earlier data indicated that the activity of PRF/GlyT2+ cells can be linked to different phases of slow cortical oscillations^23^. Here we asked to what extent the firing activity of PRF/GlyT2+ cells is sensitive to the alterations of cortical states (synchronized vs, desynchronized, Figures 4, S3A-C). We recorded PRF/GlyT2+ neurons (n=12) in GlyT2-eGFP transgenic animals and monitored the state changes in LFP activity (Figure 4A-B). In the case of 7 cells, we recorded spontaneous state changes between synchronized and desynchronized cortical states which can be observed in light ketamine-xylazine anesthesia (Figure 4B) whereas in 5 cases we evoked cortical inactivity by applying 2M potassium chloride (KCl) to the cortical surface to induce cortical spreading depression (CSD, Figure S3D-F)^32^.

**Figure 4.**
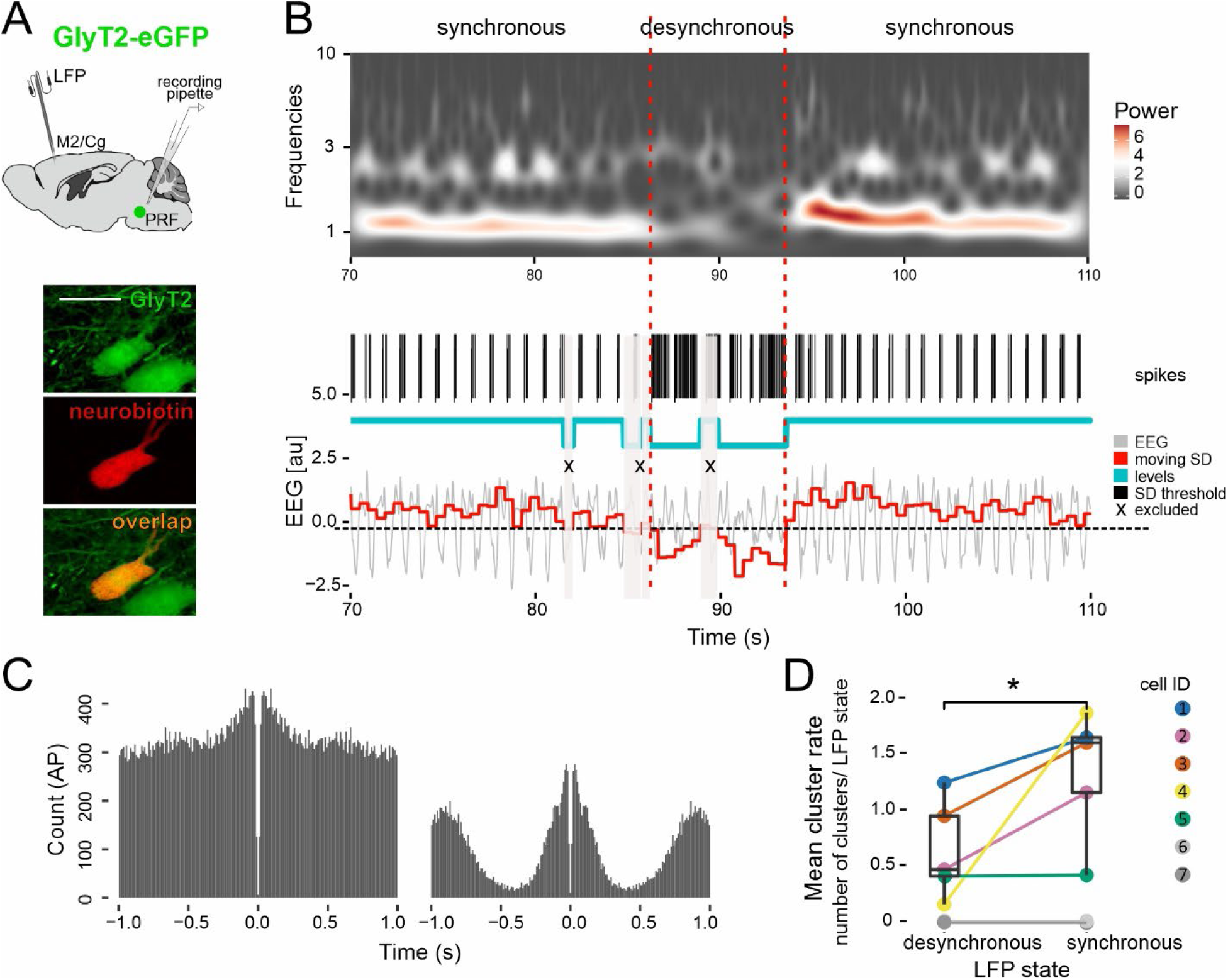
The effect of spontaneous cortical activity on PRF/GlyT2+ neurons. A) Scheme of the experiments (n= 7 mice) and a posthoc identified PRF/GlyT2+ (n=7, green) neuron filled with neurobiotin (red) and the overlapping (yellow) (bottom) B) A representative example of a spontaneous state change under light ketamine-xylazine anesthesia together with the firing activity of a PRF/GlyT2+ cell. Top: Wavelet transformation of the cortical LFP activity. Middle: Action potentials of a PRF/GlyT2+ cell, thick black lines correspond to the first action potential of the detected clusters. Blue line: synchronized and desynchronized states. Bottom: The raw LFP signal is shown in grey, and the amplitude variance of the LFP signal (moving SD in a sliding window) is in red. The black dashed line represents the variance threshold for synchronized and desynchronized periods. Periods labeled with X (gray shading) are transient states, excluded from the analysis (see Methods). C) Autocorrelogram of a representative neuron during synchronized and desynchronized periods. D) Mean cluster rate (number of clusters/LFP state length) during synchronized (n=348, mean: 1.32, SD: 0.57, median: 1.59) and desynchronized periods (n=270). The two neurons shown in gray did not fire rhythmic AP clusters. (n=7, Wilcoxon Signed-Rank Test, p=0.032) * 0.05<p; ** 0.01<p; *** p<0.001; n.s. - no significant difference. Source data are provided as a Source Data file. Scale bars: A) 20 μm.

In the first experiments, we observed that 5 out of 7 PRF/GlyT2+ cells fired rhythmic AP clusters during the slow oscillatory periods (Figures 4B-D, S3A). When the slow oscillation was replaced by a desynchronized period in the LFP, the firing pattern of the PRF/GlyT2+ cells immediately followed this transition and became irregular (Figures 4B-D, S3A). When the rhythmic cortical activity was restored, the AP clusters reappeared (Figures 4B-D, S3C). The transition from rhythmic AP cluster to tonic activity could be confirmed by autocorrelograms (Figures 4C, S3A). The mean rate of PRF/GlyT2+ action potential clusters was significantly higher in synchronized vs. desynchronized states (Figure 4D). The firing rate of PRF/GlyT2+ neurons significantly increased during the desynchronized states (Figure S3B). In the CSD experiments, cells decreased the rate of spike clusters upon cortical cell inactivation (Figure S3F). These data demonstrate a tight connection between the cortical states and firing activity of PRF/GlyT2 neurons and confirm the strong impact of the cortex on these cells during spontaneous activity.

### Co-innervation of PRF and the thalamic targets of PRF by L5 neurons

L5 neurons that innervate the striatum have been shown to arborize in the thalamus^15,16^ and since the basal ganglia also innervate the thalamus and thalamus project back to the cortex the three structures form multiple nested loops^6,8,11^.

To investigate if the same logic applies to individual L5 neurons targeting the PRF i.e. whether L5 neurons simultaneously innervate PRF and the target region of PRF/GlyT2+ cells in the thalamus (the intralaminar and parafascicular nuclei, IL/Pf) we analyzed data from the Mouse Light Neuron Browser database^33^ (https://www.janelia.org/open-science/mouselight-neuronbrowser). In the database we found 23 M2/Cg L5 neurons that had at least one axonal end point in the PRF (Figure 5A-C; Table S1.). Seventy-four percent of these neurons (n=17) projected to the IL/Pf as well. Ten out of the M2/Cg L5 17 neurons (59%) neurons co-innervating PRF and IL/Pf displayed multiple axonal end points (here defined as more than 4 end points) both in PRF and in the IL/PF. Four out of the 23 PRF projecting L5 neurons (17.4%) displayed very dense axonal arbors in PRF (more than 10 axonal end points) (Figure 5C). Three of these L5 cells innervated IL/Pf also with more than 10 axonal end points. The fourth of them formed 6 end points in IL/Pf. Confirming our anterograde tracing experiments, these neurons arborized more in the dorsomedial part of the PRF (Figure 5C). These data suggest that co-innervation of PRF and IL/Pf by individual cortical L5 neurons is common. Sixteen out of the 17 L5 neurons (94.12%) co-innervating PRF and IL/Pf also had a collateral to the striatum indicating that the cortico-PRF-thalamic message is projected to the basal ganglia as well (Table S1).

**Figure 5.**
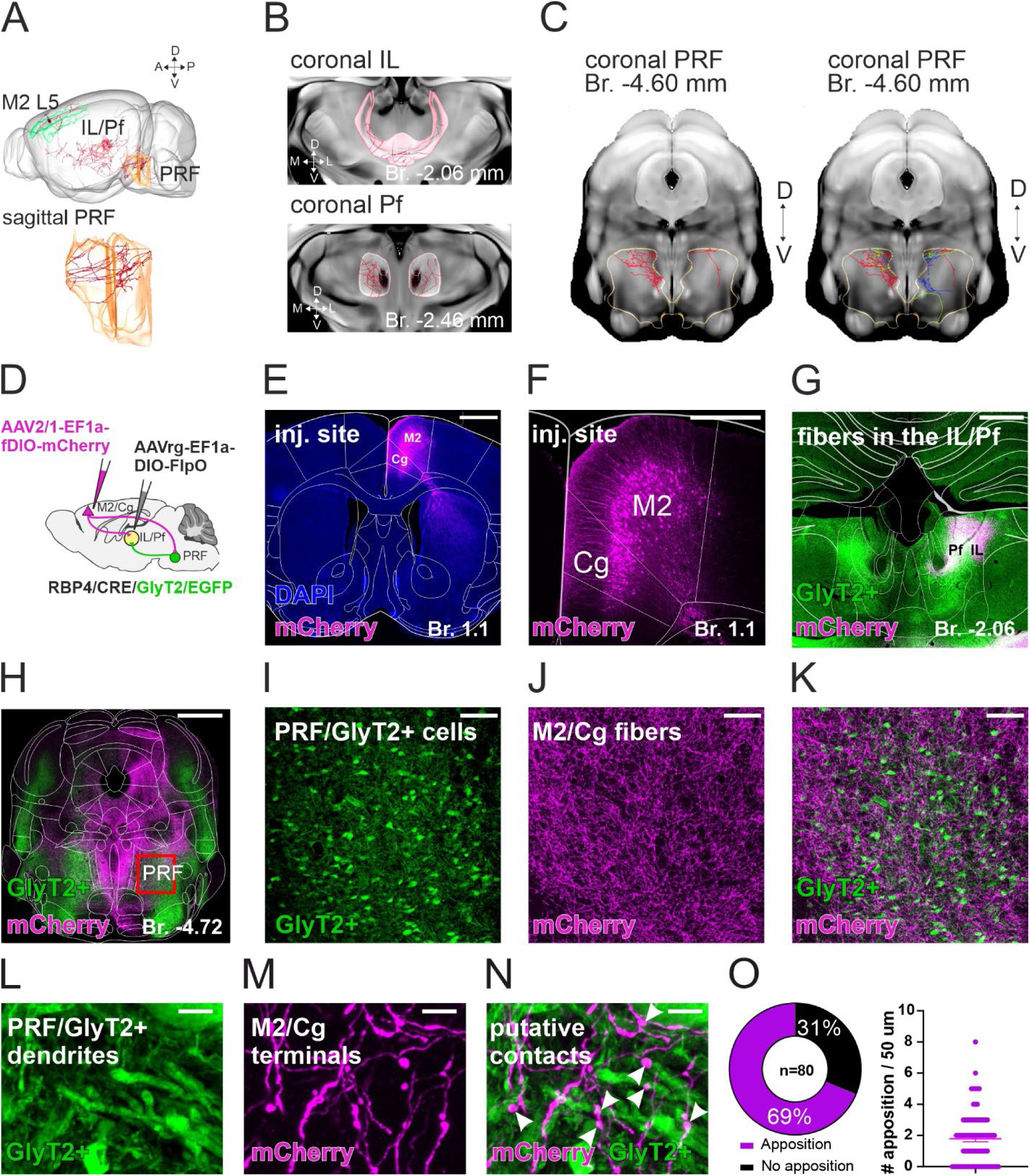
Co-innervation of PRF and thalamus by M2/Cg L5 neurons. A) Reconstruction of a representative M2 L5 (AA0245) neuron from the Mouse Light Neuron Browser database which innervates both PRF and IL/Pf (top). Axon arbor of the representative M2 L5 neuron (AA0245) at the sagittal level in the PRF (bottom). B) Axon arbor of the representative M2 L5 neuron (AA0245) in the IL (top) and Pf (bottom) C) Axon arbor of the representative M2 L5 neuron (AA0245) (left in red) and of 4 L5 neurons in the PRF (right, red, yellow, blue and green) that have 10 or more axonal end points in the PRF and also innervate IL/Pf with multiple endpoints. D) Scheme of the viral tracing to label PRF collaterals of thalamus-projecting M2/Cg cells. E) Low-power confocal micrograph of the cortical M2/Cg injection site. F) High-power confocal fluorescent image of the thalamus-projecting M2 L5 neurons. G) Merged confocal image of the labeled thalamus-projecting M2/Cg cortical fibers (magenta) and PRF/GlyT2+ fibers (green) in the thalamus. H) Merged, low power confocal image of the labeled thalamus-projecting M2/Cg cortical fibers (magenta) and PRF/GlyT2+ fibers (green) in the brainstem. I-K) Confocal micrographs of the PRF/GlyT2+ cells (green, I), anterogradely labeled thalamus-projecting M2/Cg cortical fibers (magenta, J), and their merged image (K). L-N) High power confocal microscopic image of close apposition between the PRF/GlyT2+ dendrites (green, L) and thalamus-projecting M2/Cg fibers (magenta, M). Arrows, putative contacts in (N). O) Percentage of innervated PRF/GlyT2+ dendrite (left) and the mean number of the close apposition per dendritic segment (right). Source data are provided as a Source Data file. Scale bars: E) 1 mm, F-G) 500 μm H) 1mm I-K) 100 μm L-N) 5 μm.

The Mouse Light Neuron Browser data does not permit to establish whether the cortico-PRF-thalamic neurons target PRF/GlyT2+ cells or other cell types in the PRF. In order to demonstrate that thalamic-projecting cortical cells can innervate PRF/GlyT2+ cells, we employed two experimental designs (Figures 5D and S4A). In the first configuration, we injected AAVrg-EF1a-DIO-FlpO into the IL/Pf and AAV2/1-EF1a-fDIO-mCherry into M2 in RPB4-Cre//GlyT2-eGFP mice (Figure 5D). In this case, the Flp is Cre dependent, travels retrogradely from the thalamus to the cortex, and expresses in RBP4+ L5 neurons (Figure 5E-F). The second virus is FLP-dependent and travels anterogradely from those L5 neurons which project to the IL/Pf as well (Figure 5G-K). In the second approach, we injected Cre-expressing retrograde virus (AAVrg-EF1a-Cre) into the IL/Pf and AAV5-EF1a-DIO-hChR2-mCherry virus into the M2 in GlyT2-EGFP mice (see more details in the Methods; Figure S4A).

In both experimental approaches, we observed retrogradely labeled deep layer M2/Cg pyramidal cells (Figures 5F, S4B-C) and dense axonal arbor in the dorsolateral Pf/IL verifying the injection sites and the thalamic projections (Figures 5G; S4D-H and S4M-P). Dense cortical innervation was also observed in the PRF in every case (n=4 animals), in the same PRF/GlyT2+ area (Figures 5H-K, S4I-L), which we identified after our original M2/Cg AAV5-DIO-ChR2-mCherry injections (as in Figure 1). The cortical projection was largely ipsilateral in the PRF. Using high-resolution confocal microscopy, we identified close appositions between axons of IL/Pf projecting cortical neurons and the dendrites of PRF/GlyT2+ cells (Figure 5L-N). In the first experimental configuration, 72.5% of PRF/GlyT2+ dendrites (n=40) were contacted by the axon terminals of thalamic projecting M2/Cg axons, compared to 65% in the second experiment (n=40). The mean number of close appositions per dendrite was 2.125 ± 0.308 in the first case and 1.45 ± 1.339 in the second. The pooled data show that 69% of the PRF/GlyT2+ dendritic segments (n=80) were juxtaposed to M2/Cg boutons, with an average of 1.45 ± 1.339 putative contacts per 50 µm segment (Figure 5O). Together these data show that thalamus-projecting M2/Cg pyramidal cells can also innervate PRF/GlyT2+ cells.

### Activation of PRF/GlyT2+ neurons inhibit thalamocortical neuronal activity in vivo

As the next step in the loop, we aimed to examine the impact of PRF/GlyT2+ activity on thalamic cells. PRF/GlyT2+ neurons have been shown to selectively innervate IL/Pf and evoke non-depressing, mixed, GABA/glycine receptor-mediated IPSCs on their targets^23^. However, the effect of this inhibitory input on the spike output of IL/Pf has not been investigated. To address this, we performed in vivo juxtacellular recording and labeling experiments in GlyT2-Cre mice injected with an AAV5-DIO-ChR2-eYFP virus in the PRF (Figures 6 and S5). PRF/GlyT2+ fibers were then photoactivated in the IL/Pf (Figure 6A-B). To ensure that the recorded IL/Pf neurons were located within the PRF/GlyT2+ fibers, the position of the IL/Pf cells was determined post hoc (Figure S5A). Given that IL/Pf neurons are known to be sensitive to pain^34^ we utilized urethane anesthesia to assess the impact of PRF/GlyT2+ photoactivation on spontaneous (n= 13 cells) as well as on tail-pinch-induced IL/Pf activity (n= 10 cells, (Figures 6B-C, S5B-C). The stimulation frequency used (33 Hz) was compatible with the spontaneous activity of PRF/GlyT2+ neurons measured under identical conditions^23^.

**Figure 6.**
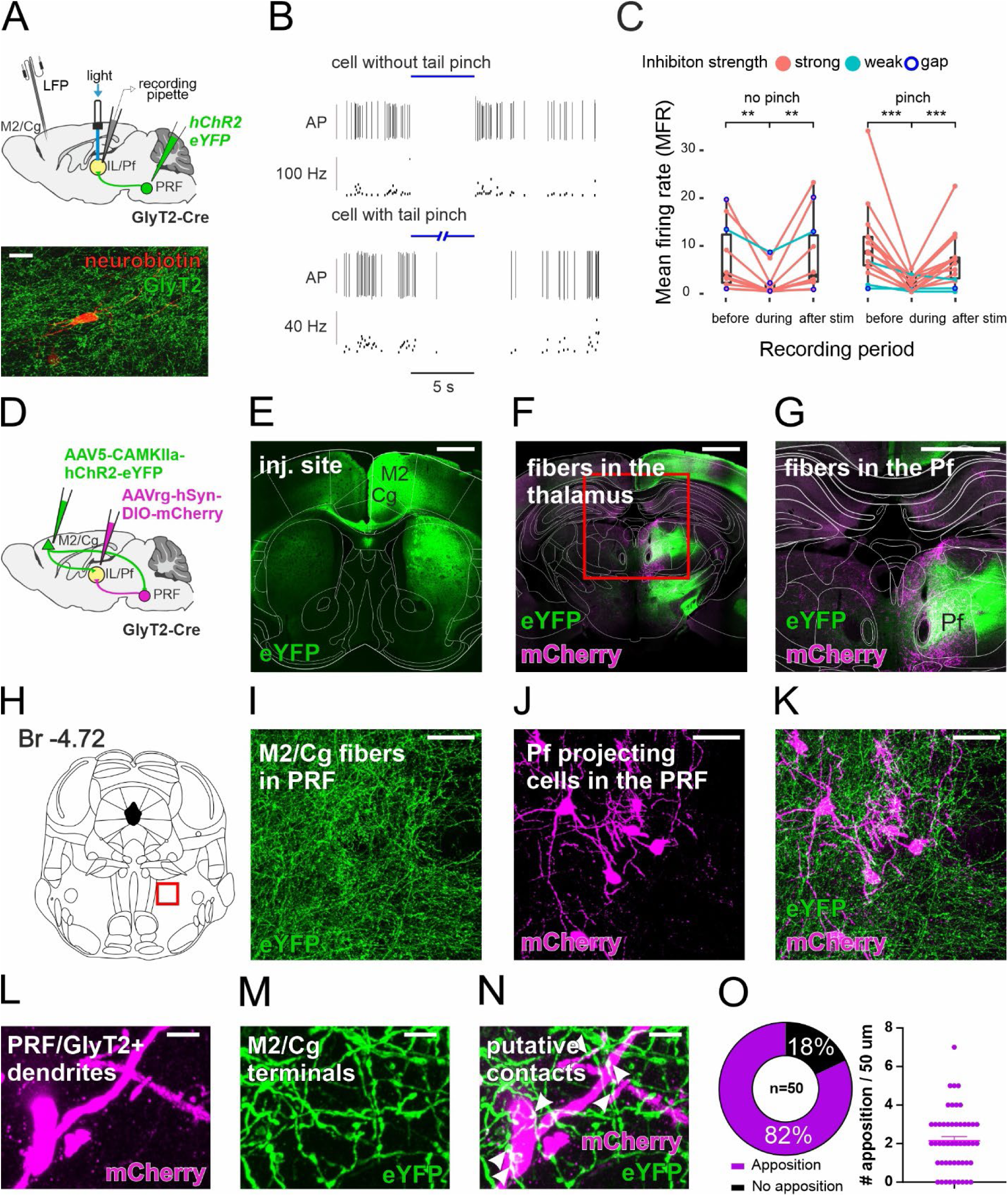
Inhibitory action and cortical innervation of thalamus-projecting PRF/GlyT2+ Cells. A) Top: Scheme of the juxtacellular recordings (n=13 mice, n=23 neurons). Bottom: Post hoc identified IL/Pf neuron surrounded by GlyT2+ fibers. B) Response of two representative IL/Pf thalamic neurons to optogenetic activation of PRF/GlyT2+ fibers (top, 5 sec stimulation, without a tail pinch, bottom, 30 sec stimulation with a tail pinch). Top: discriminated action potentials (AP), 5 sec stimulation, without tail pinch. Bottom: instantaneous firing rate (blue line = laser ON), 30 sec stimulation with a tail pinch. C) Mean firing rate (MFR) of the recorded IL/Pf thalamic neurons before, during, and after the photoactivation PRF/GlyT2+ fibers. (33 Hz, 5 ms pulse width, laser power 10 mW, 5-30 sec duration). Strongly inhibited, (red circles, n=19 cell), weakly inhibited (light blue circles, n=4, see Methods) and neurons responding with a post-stimulus gap (dark blue circle, n=4) are indicated. No pinch (left) and pinch (right) conditions are shown separately. No pinch MFR before vs. stim p=0.002; no pinch MFR stim vs after p=0.002 n=10 cells; pinch before vs. stim p=0.00012; pinch MFR stim vs. after p= 0.00085, n=14 cells; Wilcoxon Signed-Rank Test. D) Scheme of the double conditional viral tracing to label cortical inputs to thalamus projecting PRF/GlyT2+ cells. E) Low-power confocal micrograph of the cortical injection site. F-G) Merged confocal image of the M2/Cg cortical fibers (green) and PRF/GlyT2+ fibers (magenta) in the thalamus. The red rectangle in F indicates the position of the G micrograph. H) Coronal, stereotactic image of the brainstem. The red rectangle indicates the position of the I-K micrographs. I-K) Confocal micrographs of the anterogradely labeled M2/Cg cortical fibers (green, I), retrogradely labeled thalamus-projecting PRF/GlyT2+ cells (magenta, J), and their merged image (K). L-N) High power confocal microscopic image of the thalamus-projecting PRF/GlyT2+ dendrites (magenta, L) and M2/Cg fibers (magenta, M) and their putative contacts (N, arrowheads). O) Percentage of innervated thalamus-projecting PRF/GlyT2+ dendrite (left) and mean number of the close apposition per dendritic segment (right) * 0.05<p; ** 0.01<p; *** p<0.001; n.s. - no significant difference. Source data are provided as a Source Data file. Scale bars: A) 20 μm E-G) 1 mm I-K) 50 μm L-N) 5 μm.

The baseline mean firing rate (MFR) of the recorded thalamic neurons (n=23) varied between 1 and 30 Hz (Figure S5B). Tail pinch evoked a significantly higher MFR in the IL/Pf cells (Figure S5B). Optogenetic activation of PRF/GlyT2+ fibers (33 Hz) in the IL/Pf resulted in a significant decrease of IL/Pf spiking activity in both spontaneous and tail-pinch conditions at the population level (Figures 6B-C; S5C-D). The firing rate immediately returned to baseline values upon the cessation of stimulus trains (Figures 6B-C; S5C-D).

Nineteen out of the 23 IL/Pf neurons (82.6%) were considered strongly inhibited (see Methods; Figure S5E). In these neurons, inhibition of the firing output increased gradually and following a transient period of 1-2 seconds PRF/GlyT2+ inputs could almost entirely block the firing output of IL/Pf cells (Figure S5D-E). In 9 neurons we increased the duration of stimulation from 5 to 16 or 30 sec but the tonic cessation of firing was maintained for the entire duration (n=9 cell, Figure S5D). Firing activity returned to baseline levels after these long stimulations as well (Figure S5D). In the case of 4 out of the 23 IL/Pf neurons (16.7%), the PRF/GlyT2+ activation resulted in a brief gap (5-10 ms) in the PSTH immediately after the onset of the stimulus (Figure S5F-G). The time course of this precisely timed inhibition corresponded well with the kinetics of the fast GABAergic/glycinergic IPSCs recorded during the earlier in vitro observations^23^ indicating a fast, phasic inhibition in this subset of neurons. The activity of control neurons in the thalamus, located outside the PRF/GlyT2+ projection zone (n=5), was not altered by the activation of PRF/GlyT2+ fibers (Figure S5D, H).

The precise localization of IL/Pf neurons inhibited by PRF/GlyT2+ activation was verified post hoc. IL/Pf cells were localized within the PRF/GlyT2+ fibers in all cases (Figure S5A), mainly in the dorsolateral sectors. The recorded cells were distributed both in the anterior (centrolateral and paracentral) and the posterior (parafascicular) IL nuclei (Figure S5A). All weakly inhibited cells were localized in anterior IL. We conclude that PRF/GlyT2+ neurons can exert a powerful effect on the spontaneous as well as on tail pinch evoked firing activity of IL/Pf neurons and can participate in both tonic (divisive) and phasic inhibition of IL/Pf outputs.

So far, we have shown that the cortico-PRF pathway can effectively control PRF/GlyT2+ cells (Figures 3-4, S2-3) and that, in turn, PRF/GlyT2+ cells can strongly inhibit their postsynaptic partner in the thalamus (Figures 6 and S5). However, it still remains to directly demonstrate whether cortical L5 axons can actually target those PRF/GlyT2+ neurons, which innervate the thalamus and inhibit IL/Pf neurons. To investigate this question, we injected conditional retrograde AAVrg-hSyn-DIO-mCherry virus into the IL/Pf and anterograde AAV5-CAMKIIa-hChR2-eYFP into the M2/Cg of GlyT2-Cre mice (Figure 6D). We then examined the GlyT2+ cells retrogradely labeled from the thalamus in the PRF and the M2/Cg terminals contacting them. We confirmed the cortical injection sites in M2/Cg (Figure 6E) and their thalamic projection in IL/Pf (Figure 6F-G). In the brainstem, we could frequently observe retrogradely labeled PRF/GlyT2+ neurons among the cortical fibers (Figure 6H-K). Using confocal microscopy, we confirmed M2/Cg terminals formed putative contacts with 82% of all thalamus-projecting PRF/GlyT2+ dendrites (n=50, from 3 mice, Figure 6L-O). We calculated the mean number of close appositions in 50 µm long dendritic segment, which was 2.4 ± 0.22 (Figure 6O).

These data identified cortical inputs on thalamic projecting PRF/GlyT2+ cells. Together with the data showing that the same layer 5 cell can innervate PRF/GlyT2+ cells and IL/PF (Figure 5) and PRF/GlyT2+ cell contact IL/PF (Figure 6)^23^ we demonstrated that in this system cortex, PRF/GlyT2+ cells and thalamus can form multiple nested loops.

### Activation of PRF/GlyT2+ neurons promotes rotational movements

Next, we aimed to examine the behavioral effects of activating the thalamus projecting PRF/GlyT2+ cells, a key node in the cortex-PRF-thalamus communication. To address this, we injected AAV5-DIO-ChR2-eYFP into the PRF of GlyT2-Cre animals and implanted fiber optics into the thalamus (IL/Pf) (n=7 stimulus site, n=2 animals). We then photoactivated PRF/GlyT2+ fibers unilaterally and bilaterally at various laser intensities (Figure 7A) in IL/Pf. Bilateral stimulation (n=20 stimulation) or unilateral stimulation with higher (15 mW) laser intensities (n=24 stim) resulted in behavior arrest confirming earlier data^23^. Unilateral stimulation of PRF/GlyT2+ axons in the IL/Pf at lower laser intensity (1, 5 and 10 mW), however, induced rotation always in the contralateral direction (Figure 7B). Rotational movements significantly increased during stimulation and decreased after stimulus termination (Figure 7B; Video S1). Positions of the optic fibers were determined post hoc (Figure 7C).

**Figure 7.**
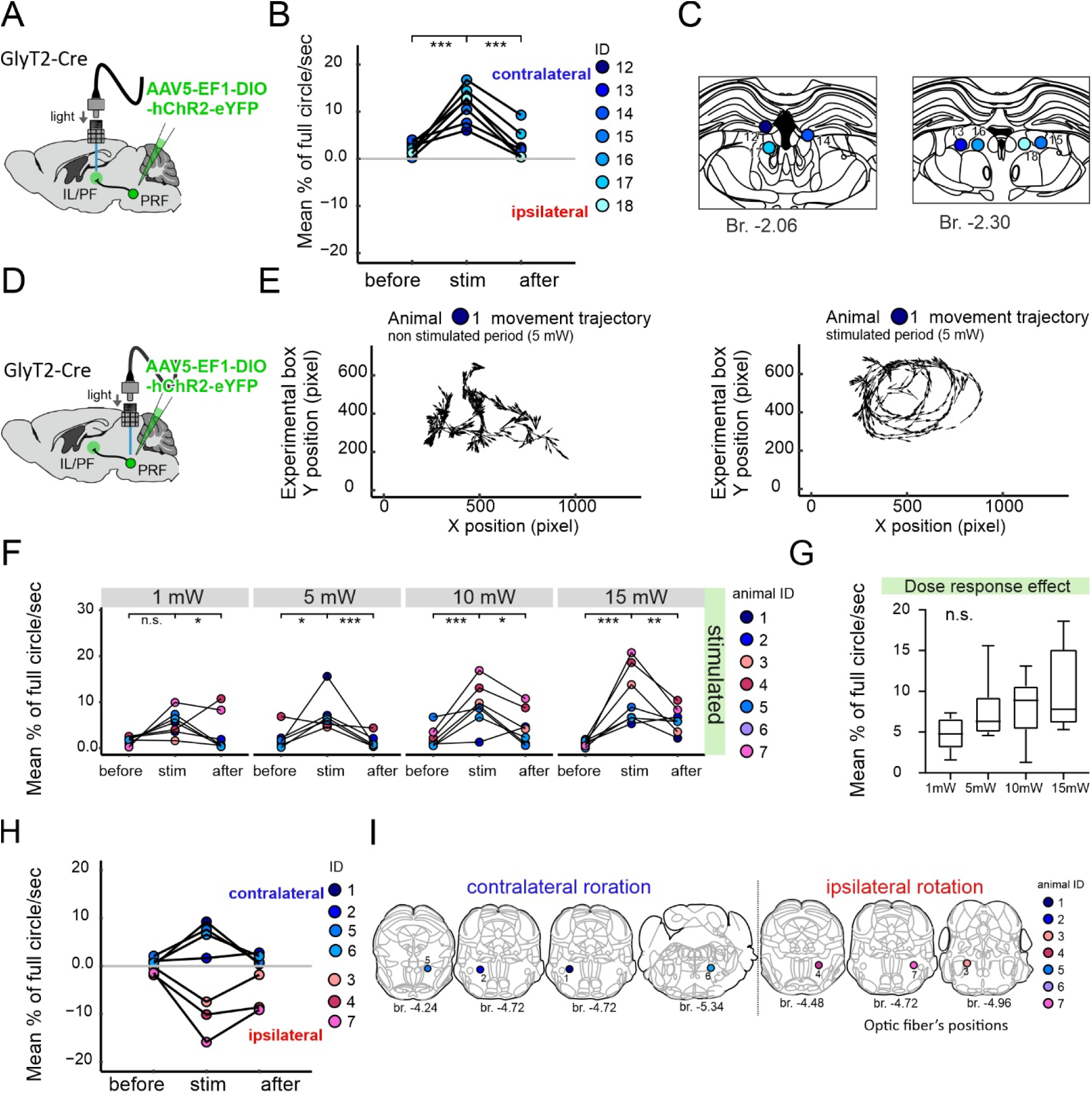
Rotational movements evoked by axonal vs somatic photoactivation of PRF/GlyT2+neurons. A) Experimental design of the photoactivation of thalamus-projecting PRF/GlyT2+ cells via their fibers in the thalamus (n=7 fiber optic fibers). B) Contralateral rotations (n=7 fiber optics) displayed as mean rotation angle before, during, and after the stimulus periods following PRF/GlyT2+ fiber stimulation in Pf. The figure represents the average percentage of a full 360° circle per second. (χ²=10.6, p=0.005, Friedman test following Durbin-Conover post-hoc, Before vs stim p<0.001; stim vs after p=<0.001) C) Schematic view of post hoc identified 7 optic fiber positions for experimental animals in Pf. D) Experimental design of the photoactivation of PRF/GlyT2+ somata in the PRF (n=7 fiber optics). E) Movement trajectory in one representative mouse displaying contralateral turning (mouse ID: 1). Non-stimulated period (left), stimulated period (right) at 5 mW in the PRF. F) Comparison of the absolute values of the mean rotation angle before, during, and after the PRF stimulus periods in the 7 experimental mice (Friedman test following Durbin-Conover post-hoc; 1 mW χ²=6, p=0.050; before vs. stim p=0.073; stim vs. after p=0,012; 5 mW χ²=8.33, p=0.016; before vs. stim p=0.038; stim vs. after p<0.001; 10 mW χ²=10.3, p=0.006; before vs. stim p<0.001; stim vs. after p=0.014; 15 mW χ²=12.3, p=0.002; before vs. stim p=<0.001; stim vs. after p=0.004 5 mW, χ²=8.333, p=0.012; G) Lack of significant dose-response effect on rotation (Friedman test χ²=7, p=0.072). H) Contralateral (n=4 animals, blueish shades) and ipsilateral (n=3 animals, reddish shades) rotations following PRF/GlyT2 somatic stimulations. The color code of the animals is the same as in F. I) Schematic view of post hoc identified optic fiber position (n=7) in PRF for the 7 experimental mice. Right: fiber optic positions inducing contralateral rotation; left: fiber optic positions inducing ipsilateral rotation. Same notation as in G). * 0.05<p; ** 0.01<p; *** p<0.001; n.s. - no significant difference. Source data are provided as a Source Data file.

These findings show that activation of thalamic projecting PRF/GlyT2+ cells via their axons in IL/Pf consistently leads to contralateral rotation. However, not all PRF/GlyT2+ cells may project to the thalamus. In order to identify, whether the behavioral effect of activating thalamus projecting PRF/GlyT2+ cells differ from that of the whole PRF/GlyT2+ population we injected AAV5-DIO-ChR2-eYFP into the PRF in GlyT2-Cre animals and implanted fiber optics, now, into the PRF and photoactivated PRF/GlyT2+ neurons unilaterally via their somata (Figure 7D) at various laser intensities (1; 5; 10; 15 mW; 5 ms pulse with, 40 Hz for 10 sec, n=7 animals).

Since the effect of PRF/GlyT2+ soma activation on movement initiation and maintenance has not been characterized first we measured the traveled distance in the first second of the PRF/GlyT2+ stimulation and the subsequent 9 seconds (Figure S6A). After the stimulus onset, we found significantly increased movement in the first second of the stimulus relative to the baseline period (before stimulus) with 3 stimulation intensities (1, 10 mW) Figure S6A). Increased locomotion persisted in the remaining 9 sec of the stimulation (Figure S6A). There was no significant dose-response effect with increasing laser light intensity (Figure S6B).

Next, we examined rotational movements during the stimulation of PRF/GlyT2+ somata (10 sec) in the same animals (Figure 7E; Video S2-3). Calculated for the whole population (n=7 optic fibers) rotational movements significantly increased compared to the baseline periods upon optogenetic activation at all stimulation intensities (1, 5, 10 and 15 mW, Figure 7F). We found no significant dose-response effect with increasing laser light intensity (Figure 7G). Interestingly, somatic stimulation of PRF/GlyT2+ neurons induced contralateral rotational movements, similar to thalamic stimulation of PRF/GlyT2+ fibers, in only 4 out of the 7 animals. In the remaining 3 animals ipsilateral rotation was observed (Figure 7H). We found no significant differences in rotational angles between the ipsi- and contralaterally turning animals (ipsi vs. contra stim for all intensity U=2 p=0.229). Post hoc histological identification of optic fiber positions within the PRF revealed no discernible anteroposterior, dorsoventral, or mediolateral differences between the animals exhibiting ipsi- or contralateral rotational movements (Figure 7I, Figure S6C-D).

In the eYFP control animals, the mean traveled distance, and the rotational angle was not significantly different between the baseline and stimulation periods (Figure S6E-K, Video S4). Next, we conducted loss-of-function experiments by injecting AAV5-DIO-eNpHR 3.0-eYFP into the PRF (n=4 animals) and implanting optic fibers into the PRF (n=8 optic fibers, Figure S7A). Similar to the gain of function experiments (Figures 7 and S6) we found that inhibition of PRF/GlyT2+ somata unilaterally could result in both contra- and ipsiversive rotations (Figure S7C, Video S5). However, for the whole group significant difference in rotational angles was not found (Figure S7C-D). Movement initiation was significantly affected only at 15mW (Figure S7E-F).

These findings demonstrate that activation of PRF/GlyT2+ neurons can elicit rotational movements. Activation of thalamic projecting PRF/GlyT2+ cells via their axons elicits contralateral turning whereas activation of the entire PRF/GlyT2+ cells population can evoke both ipsi- and contralateral turning.

### Distinct projection patterns of thalamus projecting PRF/GlyT2+ cells

Next, we examined whether the observed differences in turning behavior between PRF/GlyT2+ neurons innervating the thalamus and the entire PRF/GlyT2+ population can be explained by distinct projection patterns (Figure 8). We primarily focused on the gigantocellular nucleus in the medulla (Gi) that has recently been shown to be responsible for the motor commands of turning movements and compared ipsi- and contralateral projections^1,10,21^.

**Figure 8.**
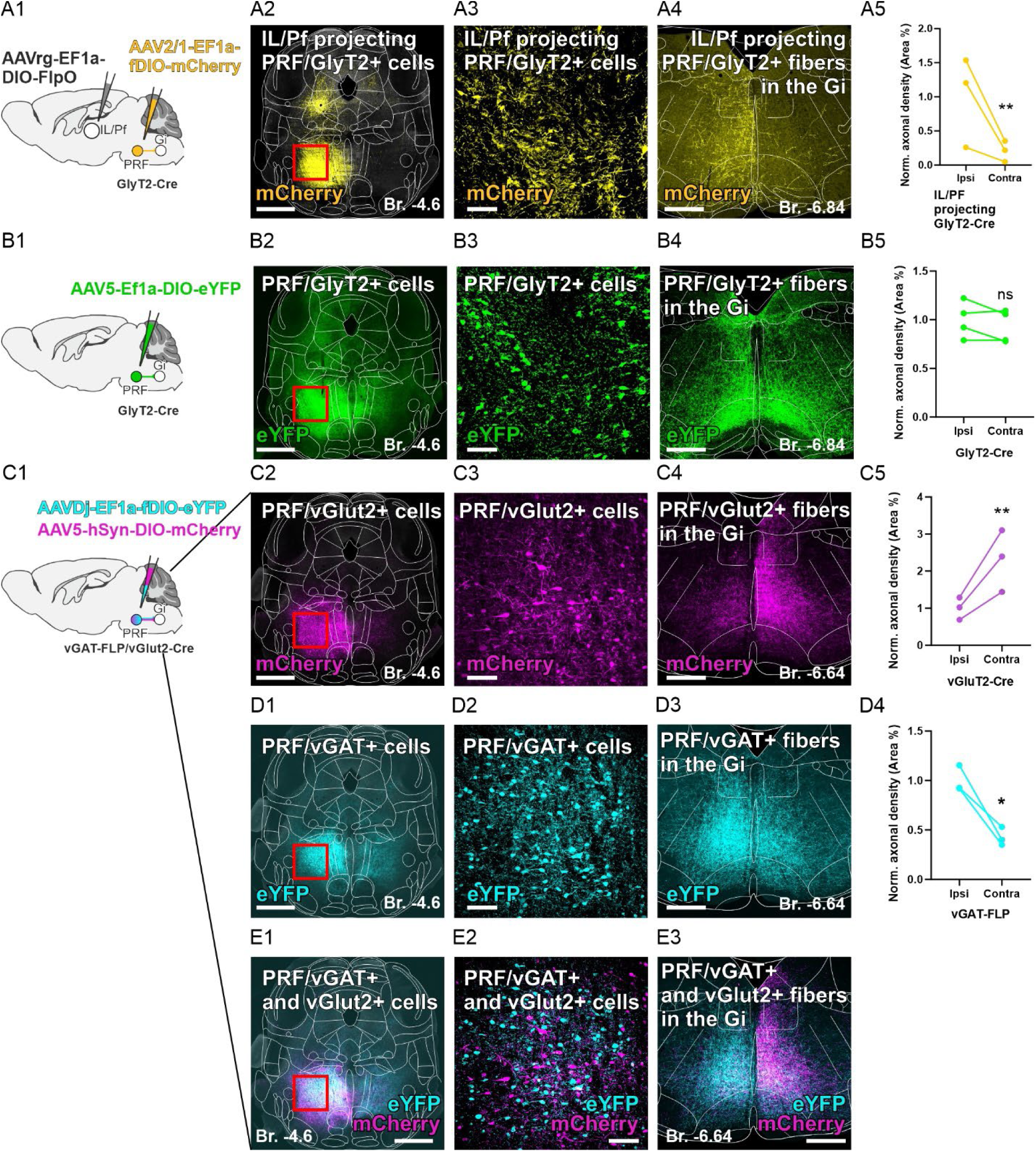
Selective ipsilateral innervation of the gigantocellular nucleus by thalamus projecting PRF/GlyT2+ cells. A1) Experimental design to label thalamus-projecting PRF/GlyT2 cells+ and their axons (n=3 mice). A2-3) Fluorescent micrograph about virus injection site in the PRF. The red rectangle on A2 indicates the position of the A3 micrograph. A4) Ipsilateral projection of the thalamus-projecting PRF/GlyT2+ cells in the Gi (yellow) A5) Quantification of ipsilateral and contralateral axonal projections of the thalamus-projecting PRF/GlyT2+ cells in the Gi in GlyT2-Cre Mice (paired t-test, t_(2)_=22.07, p=0.002, n=3 mice). B1) Experimental design to label all PRF/GlyT2 cells+ PRF (n=4 mice). B2-3) Fluorescent micrograph about virus injection site in PRF. The red rectangle on B2 indicates the position of the B3 micrograph. B4) Bilateral projection of the PRF/GlyT2+ cells in the Gi (green). B5) Quantification of the ipsilateral vs. contralateral axonal projections from PRF/GlyT2+ cells within the Gi Region in GlyT2-Cre mice (paired t-test, t_(3)_=1.462, p=0.24, n=4 mice). C1) Scheme of mixed viral injection into PRF in vGAT-Flp/vGlut2-Cre (n=3 mice). C2-3) Fluorescent micrograph about virus injection site in PRF. The red rectangle on C2 indicates the position of the C3 micrograph. C4) Bilateral projection with contralateral dominance of PRF/vGlut2+ cells in the Gi (magenta). C5) Quantification of ipsilateral and contralateral axonal projections from PRF/vGluT2+ cells within the Gi Region in vGluT2-Cre/vGAT-FLP mice (paired t-test, t_(2)_=19.04, p=0.0027, n=3 mice). D1-2) Fluorescent micrograph about virus injection site in PRF. The red rectangle on D1 indicates the position of the D2 micrograph D3) Bilateral projection with ipsilateral dominance pf PRF/vGAT cells in the Gi (cyan) D4) Quantification of ipsilateral and contralateral axonal projections from PRF/vGAT2+ cells within the Gi Region in vGluT2-Cre/vGAT-FLP mice (paired t-test, t(2)=5.61, p=0.0303, n=3 mice). E1-2) Fluorescent micrograph about virus injection site in PRF. The red rectangle on E1 indicates the position of the E2 micrograph. E3) Merged image of PRF/vGAT (cyan) and PRF/vGlut2+ (magenta) fibers in the Gi * 0.05<p; ** 0.01<p; *** p<0.001; n.s. - no significant difference. Source data are provided as a Source Data file. Scale bars: A2/B2/C2/D1/E1) 500 μm; A3/B3/C3/D2/E2) 100 μm A4/B4/C4/D3/E3) 500 μm

We investigated the output of thalamus projecting PRF/GlyT2+ cells by injecting a retrograde Cre-dependent FlpO-containing virus (AAVrg-EF1a-DIO-FlpO) into the IL/Pf thalamus and AAV2/1-EF1a-fDIO-mCherry (n=3) into the PRF in GlyT2-Cre mice (Figure 8A1). This approach selectively labels the axon arbor of thalamus-projecting PRF/GlyT2+ cells (Figures 8A, S8A-H). In order to label the axons from the entire PRF/GlyT2+ population we injected AAV5-Ef1-DIO-eYFP into PRF of GlyT2-Cre animals (n=4) (Figure 8B).

We found a stark difference in the innervation pattern of Gi between the two approaches. Thalamus projecting PRF/GlyT2+ cells provided significantly more axons to the ipsilateral Gi (Figure 8A5) whereas labeling the entire PRF/GlyT2+ population resulted in ipsi- and contralateral axons in the Gi of similar density (Figure 8B5). In addition, we found that thalamic-projecting PRF/GlyT2+ cells collateralized primarily in the ipsilateral PRF whereas we observed substantial contralateral projection after labeling the entire PRF/GlyT2+ population (Figure 8A2, B2). The thalamic projections were mainly ipsilateral in both experimental approaches as described before^23^ and targeted the dorsolateral sector of Pf (Figure S8H-I) In addition, thalamus projecting PRF/GlyT2+ cells innervated several other regions ipsilaterally including the basal forebrain, hypothalamic regions, and the PAG which are not known to be involved in turning behavior (Figure S8A-G) and are not contacted by thalamus projecting BG cells^9^. These data indicate that thalamic projecting PRF/GlyT2+ cells form a specific subpopulation of the PRF/GlyT2+ cells with a primarily ipsilateral projection pattern that is distinct from the BG outputs.

Recent data have indicated that excitatory, vGLUT2-positive PRF cells project to the contralateral Gi and are responsible for contraversive turning^10^. Our data, here, however, demonstrate that a subpopulation of inhibitory PRF cells mainly project ipsilaterally but still induce contraversive turning. In order to directly demonstrate the difference between the projection pattern of excitatory and inhibitory PRF population we utilized the fact that glycinergic cells are GABAergic as well^23,35^ and used a double transgenic vGAT-Flp//vGlut2-Cre mice to label both the excitatory and the inhibitory projections in the same animals (Figures 8C, S8J). We co-injected AAVDj-EF1a-fDIO-eYFP (cyan) and AAV5-hSyn-DIO-mCherry (magenta) unilaterally into the PRF (Figure 8C-E) and examined the Gi (n=3). We could confirm that vGlut2+ neurons provide significantly more axons to the contralateral Gi side as described before (Figure 8C), while vGAT+ neurons project bilaterally (Figure 8D), with a significantly higher fiber density on the ipsilateral side.

These experiments together confirm the presence of an inhibitory PRF cell population that projects primarily to the ipsilateral Gi and activation of these cells can have synergistic behavioral effects with the contraterally projecting excitatory PRF neurons.

## Discussion

Our data show that PRF/GlyT2+ cells are under strong cortical excitatory influence and, in turn, are able to control the spiking activity of thalamic cells in the IL/Pf. We demonstrate a loop-like organization between the cortex, PRF, and the thalamus and show how the output from this loop can influence turning behavior via the thalamus projecting PRF/GlyT2+ cells.

### Multiple nested loops

If two brain centers are connected via a reciprocal loop and in addition there is a third pathway that receives input from one of them and projects to the other, they form nested loops^11^. Here, we have shown that besides the known reciprocal connectivity between M2/Cg and IL/Pf^28,29^, i; IL/Pf-projecting M2/Cg cells target PRF/GlyT2+ cells (Figures 5 and S4), and ii; IL/Pf projecting PRF/GlyT2+ cells are among the targets of M2/Cg neurons (Figure 6). We also show that M2/Cg cells target the same sector in IL/Pf as the thalamus-projecting PRF/GlyT2+ cells. These anatomical data together demonstrate a nested loop organization in the cortex-PRF-thalamus system.

In our functional assays, M2/Cg afferents evoked strong glutamatergic synaptic responses in PRF/GlyT2+ cells both in vitro and in vivo (Figures 2-4). A particular aspect of these experiments was that high-frequency activations of the afferents were followed by faithful synaptic or spiking responses with little depression under both in vitro and in vivo conditions. In addition, alterations of spontaneous cortical activity were faithfully followed by a change in PRF/GlyT2+ firing patterns. In turn, the activation of the thalamus-projecting PRF/GlyT2+ cells via their axons in the IL/Pf exerted significant inhibitory action on their thalamic targets *in vivo* (Figure 6). Two types of responses were distinguished, a gradual, tonic reduction in firing rate (divisive inhibition) upon repetitive stimulation and the introduction of a brief gap, in the firing activity following each stimulus (phasic inhibition). The inhibited thalamic neurons were consistently found among the PRF/GlyT2+ fibers, providing morphological evidence for a direct inhibitory action. The activity of thalamic cells outside the PRF/GlyT2+ axonal arborization areas remained unaffected by the stimulation. Earlier in vitro data demonstrated non-depressing PRF/GlyT2+ IPSCs in IL/Pf cells upon repetitive PRF/GlyT2+ stimulation^23^. This synaptic action may underlie the in vivo inhibitory influence observed in this study. In summary, these data indicate that the indirect pathway between M2/Cg and IL/Pf via the PRF/GlyT2+ cells is highly effective and can modify the direct reverberating activity between the cortex and thalamus.

The cortex-PRF-thalamus loop is organized according to a similar basic logic to the cortex-BG-thalamus loop. Both systems i; link interconnected cortical and thalamic stations via an indirect inhibitory loop and ii; receive strong cortical excitation from L5 neurons which also project to the thalamus. Both BG and PRF provide powerful inhibition to their thalamic targets^23,36–39^ via large axon terminals that establish multiple synapses on a single proximal dendritic target in the thalamus^23,40^. The major difference between the two lies in the complexity of how the cortical input is processed between the input and output stations.

Previous data suggested that linking interconnected thalamic and cortical regions via an extrathalamic inhibitory transfer station which itself is under strong cortical control, maybe a common organizing principle in sensory and motor systems^27^. Both the anterior pretectal nucleus and the zona incerta have been shown to be under strong cortical influence and to exert strong inhibition on the somatosensory thalamus which itself receives powerful excitation from L5 of the primary somatosensory cortex^41–43^.

Our study demonstrated that most of the M2/Cg L5 pyramidal cells that innervated IL/Pf and PRF/GlyT2+ neurons also provided collaterals to the striatum, the input station of BG. IL/Pf is the target of both BG and PRF/GlyT2+ cells^23,29^. More specifically, a recent study found that PRF-projecting SNR cells innervate the dorso-lateral Pf/IL^9^, the same region targeted by PRF/GlyT2+ cells (this study and^23^). This indicates that despite the fact that BG and PRF/GlyT2+ cells do not interact directly^10^ they share the same cortical input pattern, and their output is integrated at the same thalamic station, IL/Pf.

Both thalamic projecting SNR and PRF neurons distribute this message to many other brain centers, but this projection pattern differs significantly. Target-specific intersectional retrograde tracings demonstrated that SNR-thalamic neurons mainly target the brainstem center involved in locomotion, orienting responses, orofacial and muscle tone control. In contrast, our study shows that the PRF/GlyT2+ cells target brain regions involved in emotional and motivational behavior, arousal, and fear (Figure S8), with the exception of ipsilateral Gi (see below). This suggests that the two loops are likely involved in different behavioral aspects of turning/orienting behavior.

### Behavioral effects

Inhibition of IL/Pf cells via the activation of thalamus-projecting PRF/GlyT2+ neurons induced contralateral turning (Figure 7). This effect is consistent with the role of IL/Pf in orienting behavior^25,26,44^ particularly with its role in primarily ipsiversive orientation^24^. However, the robust contralateral turning we observed here by activating thalamus projecting axons in the IL/Pf suggests a direct motor effect via inhibition of another brain center. Optogenetic activation of axons are known to propagate antidromically and invade axon collaterals reaching other brain regions^45^. Examination of the axonal arbor of thalamus-projecting PRF/GlyT2+ cells revealed only one output station known to be involved directly in turning behavior, the Gi (Figures 8 and S8). The presence of these descending collaterals confirmed earlier observations on a PRF-Gi pattway^46,47^. Our study specifically demonstrated that while the entire PRF/GlyT2+ cell population projects both to the ipsi- and contralateral Gi equally, the thalamic projecting subpopulation exhibits a very strong bias toward the ipsilateral Gi. Importantly, it is known that the inhibition of the reticulospinal, Chx10-expressing Gi neurons results in contralateral turning whereas their activation has opposite effect^20,21^. Thalamic projecting PRF/GlyT2+ neurons have a dual, GABAergic, glycinergic phenotype and, as we show here, they can potently inhibit their targets (Figures 6 and S5). Based on these we propose here that the contralateral turning after the activation of thalamic projecting PRF/GlyT2+ cells arises primarily via the inhibition of ipsilateral Gi through their descending collaterals. This is confirmed by the fact that during contralateral turning, ipsilateral Gi cells are inhibited and display a strong transient reduction in their activity^10^. Thus, based on our results, the contralateral turning induced by activating thalamus-projecting inhibitory PRF/GlyT2+ cells is synergistic with the activation of excitatory PRF/vGLUT2+ cells^10^. In contrast to thalamus-projecting PRF/GlyT2+ cells, PRF/vGLUT2+ cells project to the contralateral Gi. Still their activation induce contralateral turning via the activation of contralateral Chx10 expressing neurons (Fig 8 and ^10^). Therefore, activation of PRF can lead to contralateral turning both via a descending, ipsilateral, inhibitory pathway and a descending, contralateral excitatory pathway to Gi.

As our data show, the activation of PRF may arise from the cortex and indeed M2/Cg cortex has already been implicated in orchestrating orienting behavior ^48–50^.

In contrast to photoactivation of thalamus-projecting PRF/GlyT2 cells contralateral turning could be elicited in only 4 out of the 7 animals following the stimulation of PRF/GlyT2+ cell bodies. In 3 animals with PRF/GlyT2+ cell body stimulation, ipsilateral, not contralateral rotation occurred suggesting a different neuronal mechanism. Importantly, we did not find systematic differences in the localization of the optic fibers inducing either ipsi- or contralateral rotation. In parallel with these behavioral data, when we labeled the descending axons of the entire PRF/GlyT2+ cell population to Gi we found no difference between the innervation of ipsi- and contralateral sides. In contrast, there was a very strong ipsilateral bias when we were labeling the thalamus-projecting PRF/GlyT2+ cells (see above). This suggests the existence of a sizable non-thalamus-projecting PRF/GlyT2+ cell population that innervates the contralateral side rather than the ipsilateral side. Activating these cells would inhibit contralateral Gi, obviously leading to ipsilateral turning. Thus, our data suggest that circuit elements responsible for ipsi- or contralateral rotation in the PRF are intermingled. These results align well with recent discoveries demonstrating the modularity or granularity of basal ganglia and brainstem motor networks^2^. According to these findings, neuronal populations with specialized functions in the brainstem can be located in close proximity and still coordinate different types of motor program. Along these lines, it has been previously shown that in the PRF, stimulation sites including unilateral or bilateral atonia via the spinal cord are mixed in the PRF^19^, indicating that granularity^2^ may also apply to this region.

### Open questions

Our study leaves several questions open for future investigations. First, many cortical inputs were not in close apposition to PRF/GlyT2+ cells suggesting the PRF/vGLUT2+ neurons might also be contacted. However, the direct synaptic input and action of cortex on vGLUT2+ neurons remain to be established. Second, in parallel with inhibitory PRF cells, vGLUT2+ PRF cells also provided input to the same sector of IL/Pf (Figure S8). The role of this “push-pull” projection from PRF to the thalamus requires further investigation. Finally, in contrast to the PRF/GlyT2+ labeling that highlighted similar innervation of both sides of Gi equally, tracing PRF/vGAT+ cells revealed a strong ipsilateral bias in Gi (Figure 8D). Since all GlyT2+ cells are vGAT+ as well^23^ this finding suggests the presence of a sizable vGAT+/GlyT2-cell population in the PRF that provides strong ipsilateral inhibition and thus can participate in contralateral turning.

### Conclusions

Our data show that cortical message arising from higher order motor cortices can be processed parallelly by two distinct cortico-subcortical loops: the cortex-BG-thalamus and the cortex-PRF-thalamus loops. The two systems have distinct output to the brainstem still can have synergistic effects on turning behavior by targeting distinct neuron populations in the PRF.

## STAR Methods

### Animal housing

All experimental procedures were approved by the Institutional Ethical Codex, Hungarian Act of Animal Care and Experimentation (1998, XXVIII, section 243/1998) and the Institutional Animal Care and Use Committee of the Institute of Experimental Medicine, Hungarian Research Network, Budapest and by the regulations of the European Union guidelines (directive 2010/63/EU). The experiments were performed by the National Animal Research Authorities of Hungary (PE/EA/877-7/2020). We used healthy, adult male mice (40-150 days) for the experiments. We used the following mice strains: GlyT2-Cre (Tg) - B6129F1 and Bl6Fx, Rbp4/cre//BAC_glyt2/GFP (TgTg) - C57Bl/6J, vGAT/t2A-Flpo//BAC-vglut2/icre (TmTg)-C57Bl/6J, BAC_glyt2/GFP -C57Bl/6J^29^. Mice were entrained in a 12-hour light/dark cycle with food and water available as *ad libitum*. We performed animal research according to the 3R principles.

### Surgeries

#### Anesthesia

The surgeries were done for all experiments (anatomy, electrophysiology, optogenetics) based on the following protocol. As anesthetics, mice received an intraperitoneal injection of ketamine (111 mg/kg, Produlab Pharma, #07/01/2302) and xylazine (4.3 mg/kg, Produlab Pharma, #07/03/2303). For the maintenance of the anesthesia, intramuscular injection of ketamine/xylazine was given every 30-50 min during the experiments. The head of the animal was fixed in a stereotaxic apparatus (David Kopf Instruments, Tujunga, California 91042, Model 900 Small Animal Stereotaxic Instrument). In one experimental condition, we used urethane anesthesia during IL/Pf in vivo juxtacellular recordings (0.12–0.15 g/100 g).

#### Virus injections and fiber optics implantations

For the anatomical investigations and *in vitro* and *in vivo* electrophysiological experiments, all stereotaxic tracer injections were performed using a glass pipette (intraMARK, 20-30-μm tip diameter, BLAUBRAND, injection flow: 25nl/min) connected to a syringe and a stereotaxic micromanipulator (Kopf Instruments). After injection, the capillary was left at the injection site for 5-10 min before slow withdrawal to allow diffusion and minimize backflow. The mice were perfused or were used for experiments following a 3–6-week survival time.

For the investigation of the cortico L5-PRF pathway, we used Rbp4/cre//BAC_glyt2/GFP (TgTg) mouse line (altogether n=25 mice)(n=11 for the anatomical investigation, n=7 for *in vitro* electrophysiology and n=7 for *in vivo* electrophysiology)

For all (in vivo and in vitro) electrophysiology and nine of the anatomical experiments the following virus injection design was used. We injected AAV5.EF1.dflox.hChR2(H134R)-mCherry.WPRE.hGH (based on Addgene plasmid #20297, UNC Vector Core) cre-dependent virus into the frontal cortex, M2/Cg (AP: Br.1 mm and 2 mm; ML: Br.0.5 mm; DV: 0.7 mm; bilaterally into 2-2 AP position, 80 nl each, 4 injection/mouse). For *in vivo* electrophysiology L5 photoactivation, the optic fibers (100 µm, 0.22 NA) were positioned above the virus injection site (M2/Cg).

For labeling the axon collaterals of PF targeting L5 neurons we used two types of virus injection design (n=4), where in two caseswe injected AAVrg-EF1a-DIO-FlpO-WPRE-HGHpA (Addgene #87306-AAVrg) virus into the Pf unilaterally (AP: −2, ML: 0,5, DV: 3; 30 nl). After three weeks we injected AAV2/1-EF1a-fDIO-mCherry (Addgene #114471-AAV2) virus into the M2/Cg (AP: Br.1 mm and 2 mm; ML: Br.0.5 mm; DV: 0.7 mm; unilaterally into 2 AP position, 80 nl each, 2 injection/mouse).

The other two cases we used <u>BAC_glyt2/gfp</u> mouse line (n=2) and we injected AAVrg-EF1a-Cre (Addgene # 55636-AAVrg) into the Pf unilaterally (AP: −2, ML: 0,5, DV: 3; 30 nl). We waited three weeks and injected AAV5.EF1.dflox.hChR2(H134R)-mCherry.WPRE.hGH (based on Addgene plasmid #20297, UNC Vector Core) into the M2 (AP: Br.1 mm and 2 mm; ML: Br.0.5 mm; DV: 0.7 mm; unilaterally into 2 AP position, 80 nl each, 2 injection/mouse).

The last experimental design of the cortico L5-PRF pathway’s investigation was labeling M2 axons on Pf-projecting PRF/GlyT+ neurons in GlyT2-Cre mice (n=4). We injected AAV5-CAMKIIa-CHR2(H134R)-EYFP (Addgene #26969-AAV5) into the M2 (AP: Br.1 mm and 2 mm; ML: Br.0.5 mm; DV: 0.7 mm; unilaterally into 2 AP position, 80 nl each, 2 injection/mouse) and AAVrg-hSyn-DIO-mCherry (Addgene #50459-AAVrg) into Pf unilaterally (AP: −2, ML: 0,5, DV: 3; 30 nl).

For the investigation of the PRF/GlyT2+-thalamic pathways altogether n=32 adult GlyT2-Cre male mice were used including the in vivo optogenetic behavioral (n=17),electrophysiology (juxtacellular recording, n=13) experiments and anatomical investigation (n=3). For optogenetic activation of PRF/GlyT2+ cells, we injected AAV5.EF1a.DIO.hChR2(H134R)-eYFP.WPRE.hGH (based on Addgene plasmid #20298, UNC Vector Core).For optogenetic activation of PRF/GlyT2+ fibers in the IL/Pf we did the same injections but bilaterally. For optogenetic inhibiton of the PRF/GlyT2+ somata, we injected AAV5-Ef1a-DIO eNpHR 3.0-EYFP (Addgene #26966-AAV5) into PRF bilaterally (AP:-4.4, ML:0.8, DV: 4.2, 100-100 nl). In the control experiment we injected AAV5.EF1a.DIO.eYFP.WPRE.hGH (based on Addgene plasmid #27056, Penn Core) virus into the PRF unilaterally (AP:-4.4, ML:0.8, DV: 4.2, 100 nl).

After virus injection the optic fibers (Thorlabs, FG105UCA, Ø105 μm core, 0.22 NA) were implanted into the PRF (n=15 optic fiber in n=11 mice, AP:-4.4, ML:0.8, DV: 4.2) or IL/PF (n=7 optic fiber in n=2 mice, AP:-1.9 or 2.3 mm; ML: −0.8 mm; DV: −2.5 or −2.8 mm). Mice recovered from surgery and the viruses could transfect the PRF/GlyT2+ cells for 3 weeks before behavioral testing or juxtacellular recording. The optic fibers were labeled with DiI stain (1,1’-Dioctadecyl-3,3,3’,3’-Tetramethylindocarbocyanine Perchlorate (’DiI’; DiIC18(3); ThermoFisher Scientific, #D282).

For further investigation of the PRF/GlyT2+-thalamic pathways, we labeled the axon arbors of PF targeting PRF/GlyT2+ cells (n=3 in GlyT2-Cre mice). We injected AAVrg-EF1a-DIO-FlpO-WPRE-HGHpA (Addgene #87306-AAVrg) virus into the Pf unilaterally (AP: −2, ML: 0,5, DV: 3; 30 nl). After three weeks we injected AAV1-EF1a-fDIO-EYFP (Addgene #55641-AAV1) or AAV1-Ef1a-fDIO mCherry (Addgene #114471-AAV1) into PRF unilaterally (AP:-4.4, ML:0.8, DV: 4.2, 100 nl).

For examining PRF/vGluT2+ and PRF/vGAT+ projections in the Gi we used vGAT/t2A-Flpo//BAC-vglut2/icre mice (n=3). We injected mixed AAV5-hSyn-DIO-mCherry (Addgene #50459-AAV5) and AAVDJ-EF1a-fDIO-EYFP-WPRE viruses (based on Addgene plasmid #55641, UNC Vector Core) into the PRF unilaterally (AP:-4.4, ML:0.8, DV: 4.2, 100 nl; 1:1 ratio of the viruses).

### Histology

#### Perfusion and tissue fixation

Perfusions were performed under deep anesthesia (sodium pentobarbital 60 mg/kg). Using saline and a fixative solution (0.1% glutaraldehyde (wt/vol) (Electron Microscopy Sciences, #16210) and 4% paraformaldehyde (wt/vol) (TAAB Laboratory, #P001) in 0.1 PB. A peristaltic pump perfused the solutions through the ascending aorta. After the fixation, the brain was sectioned using a vibratome (50 µm coronal slices).

#### Immunohistochemistry

For all immunostaining methods, we used TBS buffer. The sections were incubated in sucrose solutions for cryoprotection and were frozen above liquid nitrogen to help the penetration of the antibodies, followed by BSA treatment (4% 40 min). To visualize the L5 fibers (ChR2-mCherry viral tracer) we used a rabbit anti-mCherry primary antibody (1:4000 or 1:1000, BioVision, Inc., 23 California 95035, #5993-100, overnight). To label GlyT2+ cells and fibers in the PRF and in the IL/Pf (in the case of eYFP containing viral tracers or in the Rbp4/cre//BAC_glyt2/GFP mouse line), the sections were treated with chicken anti-eGFP primary antibody (1:15000 or 1:1000, ThermoFisher, #A10262).

For light and electron microscopic analysis when we visualized the L5 fibers in the brainstem after the anti-mCherry primary antibody we used anti-rabbit ImmPRESS (1:2, Vector Laboratories Burlingame, Ca 94010) and DAB-Ni/DAB (DAB-Ni: Nickel-intensified 3,3′-diaminobenzidine (bluish-black reaction product), DAB: 3,3′-diaminobenzidine, Sigma-Aldrich, #D5637, brown reaction product) as a chromogen. In the other case, after the anti-eGFP primary antibody (1:15000, ThermoFisher, #A10262) we used ABC (avidin biotinylated horseradish peroxidase complex, 1:300, Vector Laboratories, #PK-4000) and DAB as a chromogen.

For fluorescent and confocal microscopic investigations of L5 fibers, after the anti-mCherry primary antibody, we used Cy3-conjugated goat anti-rabbit secondary antibody (1:500, Jackson Immunoresearch, RRID: AB_2313593; Code: 111-167-003) or Cy3-conjugated donkey anti-rabbit (1:500, Jackson Immunoresearch, RRID: AB_2307443; Code: 711-165-152). In the case of the anti-eGFP primary antibody, we used the anti-chicken-Alexa488 secondary antibody (1:500, ThermoFisher, #A-11039) or Alexa 488 donkey anti-chicken secondary antibody (1:500, Jackson Immunoresearch, RRID: AB_2340375; Code: 703-545-155) to investigate GlyT2//eGFP components.

After juxtacellular recordings neurobiotin was visualized with Cy3-conjugated streptavidin (1:2000, Jackson Immunoresearch RRID: AB_2337244; Code: 016-160-084) in the PRF or in the IL/Pf. The GlyT2+ neurons/fibers and the cortical axons were visualized using the same histology mentioned above so we developed DAB-Ni/DAB or fluorescent immunostaining respectively.

To localize the fiber optics and juxtacellularly labeled cells, check virus injections, and analyze the fibers in the PRF, all micrographs were taken with OLYMPUS BX61, FLUOVIEW FV1000 confocal microscopy (software: Olympus Fluoview 1.6) or with ZEISS AxioPlan2 fluorescent microscopy with OLYMPUS DP70 camera (software: Olympus DPController 1.2.1.108) or Zeiss Axio Imager M1 microscope coupled to an AxioCam HrC digital camera.

#### Fluorescent microscopy

To verify virus expression and optic fiber placement, fluorescent images were captured using a Panoramic Digital Slide Scanner (Zeiss, Plan-Apochromat 10X/NA 0.45, xy: 0.65 μm/pixel, Panoramic MIDI II; 3DHISTECH, Budapest, Hungary). Every sixth section was stained and analyzed for each animal. Axonal projections in the Gi originating from different cell populations in the PRF were quantified by delineating regions of interest (ROIs) in the Gi (ipsilateral and contralateral, each 0.25 mm²) based on neuroanatomical landmarks identified using Hoechst counterstaining on fluorescent images, which were delineated with SlideViewer software (3DHISTECH).

#### Axonal density analysis

For intensity density analysis, FIJI was used to split the color channels of each image into red, green, and blue. The relevant channel (red for RFP, green for eGFP) was converted to grayscale. Background noise was removed, and the 8-bit images were then converted to binary with a consistent threshold for the reporter protein (RFP or eGFP). This allowed for precise calculation of axonal density within the selected regions. Axonal density in each image was calculated and expressed as the mean area percentage. To compare the ipsilateral and contralateral axonal densities statistically, the data were normalized to the ipsilateral side.

#### Density mapping analysis

Following fluorescent immunostaining, we investigated the density of L5 fibers in the PRF. A map based on fiber density was created by FIJI software16^62^. A high-magnification confocal micrograph was taken about PRF/GlyT2+ cells and M2/Cg L5 fibers (20x, NA: 0.75). The pictures were converted to 8-bit in depth then the brightest pixels of single optical slices of the original image stacks were projected into one plane to depict all of the labeled bright structures. These maximum intensity projections were smoothed by a Gauss filter with a radius of 40 pixels then the pixels were ordered according to their intensity values in 8 categories. The result is a pseudocolored, semi-quantitative map of the fiber density that enables the localization of the highest fiber density areas in a given image.

#### Confocal microscopy

Confocal images of the M2/Cg1, IL/Pf, and PRF regions were captured using a Nikon C2 Confocal Laser Scanning Microscope. The imaging was performed with a 4x Plan Fluor objective (NA 0.13, xy resolution 2 µm/pixel), a 10x Plan Fluor objective (NA 0.13, xy resolution 0.63 µm/pixel, z-step size 2 µm), and a 20x CFI Plan Apo VC objective (NA 0.75, xy resolution 0.32 µm/pixel, z-step size 1 µm, Nikon Europe). Confocal microscopy was used to analyze the spatial relationship between M2/Cg1 axonal boutons and labelled dendritic segments in the PRF. eYFP fluorescence was amplified using anti-GFP labelling, and for close apposition analysis, RFP fluorescence was amplified with anti-RFP labelling. To assess close appositions between axonal boutons and dendritic segments in the PRF, high-resolution z-stack images (Nikon CFI Plan Apo VC60X/NA 1.40 Oil objective, 0.09 µm/pixel, z-step size 0.3 µm) were taken from 5 µm of the slice. Confocal images were acquired with consistent settings for pinhole size, gain level, axial section depth, and laser intensity across all PRF slices. Close appositions, defined as potential synaptic contacts between labeled axonal boutons and dendritic segments, were manually quantified using NIS Elements Software. The analysis focused on identifying and counting instances where eYFP- or RFP-labeled boutons were in close proximity to EGFP- or RFP-labeled dendritic segments, indicating possible synaptic connections.

#### Electron microscopy

To analyze the M2/Cg terminals’ ultrastructure, we performed immunostaining for the ChR2-mCherry viral tracer (DAB, as described above). To confirm the innervation of PRF/GlyT2+ neurons from M2/Cg, we performed double immunostaining for the ChR2-mCherry viral tracer (DAB-Ni) and GlyT2+//eGFP (DAB) and processed the material for EM. In our hands, the modified DAB-Ni precipitate in the axon terminals could be unambiguously differentiated from the end product of marble-like DAB.

After double immunohistochemistry of L5 terminals and PRF/GlyT2+ cells the sections were treated with OsO_4_ (0.5%, vol/vol, with 7% sucrose for 40 min in 0.1 M PBS), followed by dehydration, sequential incubation in ethanol, uranile acetate, and acetonitrile for ultrastructural analysis. The dehydrated sections were embedded in Durcupan (Sigma Aldricht, #44610Aldrich). Blocks containing PRF were re-embedded and sectioned to ultrathin sections (60 nm thick) with an Ultramicrotome (EM UC6, Leica Biosystems). The dendrites and terminals (area, diameter) were measured in three non-consecutive sections in FIJI software^62^.

The sections were examined using a HITACHI 7100 electron microscope; the electron micrographs were taken with a Megaview digital camera.

### In vitro electrophysiology

The *in vitro* electrophysiological experiments of Figure 2 were performed on coronal slices from the brains of PRF of adult mice (more than 3 months old), see the virus injection method in the previous paragraph.

Slices were prepared for a minimum of 4 weeks following surgery. Briefly, mice were anesthetized with an intraperitoneal injection of pentobarbital, then perfused through the ascending aorta with cold (2°-5°C) bicarbonate-buffered saline (BBS) containing (in mM): 115 NaCl, 2.5 KCl, 1.6 CaCl_2_, 1.5 MgCl_2_, 1.25 NaH_2_PO_4_, 26 NaHCO_3_ and 30 glucose. Mice were then decapitated, and the portion of the brain containing the PRF was extracted and placed in BBS at 2°-5°C for a few minutes. Slices (300 μm) were then cut with a ceramic blade on a 7000smz-2 vibratome (Campden). The slicing procedure was performed in an ice-cold solution containing (in mM): 130 potassium gluconate, 15 KCl, 0.05 EGTA, 20 Hepes, 25 glucose, 1 CaCl_2_ and 6 MgCl_2_ supplemented with D-APV (50 µM). Slices were then transferred for a few minutes to a solution containing (in mM): 225 d-mannitol, 2.5 KCl, 1.25 NaH_2_PO_4_, 25 NaHCO_3_, 25 glucose, 1 CaCl_2_ and 6 MgCl_2_, and finally stored for the rest of the experimental day at 32°-34° C in oxygenated BBS (pH 7.4 after equilibration with 95 % O_2_ and 5 % CO_2_). For all recordings, slices were continuously superfused at 32°–34° C with oxygenated BBS.

We recorded GFP-positive neurons from the pontine reticular formation (PRF; ^29^). PRF identification was facilitated by the presence of the red fluorescent plexus originating from the cortical viral injections, observed via a 530 nm green LED (Thorlabs). Cells were visualized with a combination of infrared Dodt contrast and an on-line video contrast enhancement, using a CoolSnap HQ2 CCD camera (Photometrics) run by MetaMorph (Universal Imaging). Fluorescent glycinergic neurons were identified using a 470 nm-wavelength blue LED (Thorlabs). Both 470 nm and 530 nm LEDs were coupled to the slice chamber via the epifluorescence pathway of the microscope.

Whole-cell, voltage clamp recordings were performed with an EPC-10 double amplifier (Heka Elektronik) run by PatchMaster software (Heka). Patch pipettes (resistance 2-3 MΩ) were filled with an intracellular solution containing (mM): 110 CsMeSO_3_ (Sigma), 4.6 MgCl_2_, 10 HEPES, 10 K_2_-creatine phosphate (Calbiochem), 10 TEA-Cl (Sigma), 0.05 4-AP (Sigma), 4 Na_2_-ATP, 0.4 Na_2_-GTP, 1 QX-314 (Tocris), pH 7.35 with CsOH (∼300 mOsm). Series resistance was partially compensated (max 65%), whereas liquid junction potentials were not corrected. Stimulation train recordings were performed at a holding potential of −60 mV. The AMPA and NMDA components of the EPSCs were examined at both −60 mV, and +50 mV. The brief (2–3 ms long) flashes of blue light used to evoke synaptic responses were provided by triggering the 470 nm LED. TEA-Cl (500 µM) and 4-AP (50 µM) were added to the perfusing BBS when the AMPA and NMDA components were investigated. This procedure was necessary both for increasing synaptic release probability, and for reducing the sizeable current leaks developing at +50 mV in control recording conditions, thus decisively improving EPSC detection.

### In vivo electrophysiology

#### Cortical activation and LFP recording

For in vivo investigation of the L5-PRF pathway we activated the M2/Cg L5 cells optogenetically (see above the surgery details in Surgeriess n=7) and electrically (in BAC_Glyt2/GFP mice, n=3). The laser beam was generated by a 473 nm laser (Thorlabs lasers). The laser pulses were 5 ms long, 10 mW intensity, and were delivered at 1,10 and 20 Hz frequencies (n=50 stimulus in 5 trains). When we applied electrical stimulation (1-2 μA) we used a bipolar stimulating electrode on the surface of the cortex (M2/Cg, same position described above).

To record cortical field potentials, bipolar LFP electrodes (FHC, resistance ∼1 MΩ) were inserted into the frontal cortex of mice (Bregma 1.7 mm; lateral −0.8 mm). The recorded signal was amplified, band-pass filtered from 0.16 Hz to 5 kHz for LFP recordings and from 100 Hz to 5 kHz to record the multiunit activity (Supertech BioAmp, Supertech, Pécs, Hungary) and digitized at 20 kHz (micro 1401 MkII, CED, Cambridge, UK).

#### Juxtacellular recording

PRF and IL/Pf single unit activity was recorded via glass microelectrodes (in vivo impedance of 20-40 MΩ) made from borosilicate glass capillaries (1.5 mm outer diameter, 0.75 or 0.86 inner diameters, Sutter Instrument Co., Novato, CA, USA or WPI Inc. Sarasota, Fl, USA). Electrodes were positioned into the PRF (Bregma −4.4 mm; lateral −0.8 to −1 mm; ventral from cortical surface −3.8 to −4.8 mm) or IL/Pf (Bregma −1.9 to 2.3 mm; lateral −0.8 mm; ventral from cortical surface −2.5 to −2.8 mm) (using a piezoelectric microdrive (Burleigh 6000 ULN or ISS 8200, EXFO, Quebec City, Quebec, Canada). Neuronal signals were amplified by a DC amplifier (Axoclamp 2B, Axon Instruments/Molecular Devices, Sunnyvale, CA, USA), further amplified, and filtered between 0.16 Hz and 5 kHz by a signal conditioner (LinearAmp, Supertech) and recorded by Spike2 5.0 (CED). The electrodes were filled with 0.5 M K+-acetate and 2% neurobiotin (Vector Laboratories, Burlingame, CA, USA, #SP-1120). Juxtacellular labeling of the recorded neurons was done by filling the neurons with neurobiotin, as described in^37^. After the experiments, the animal was perfused, and coronal sections were cut from the whole brain (3.1. Perfusion and tissue fixation). To visualize neurons filled with neurobiotin and to determine their GlyT2 content (in the PRF) or whether it is surrounded by PRF/GlyT2+ fibers (in IL/Pf), we treated the sections as described in 3.2. Immunohistochemistry. The GlyT2::eGFP positivity was determined by confocal microscopy.

#### Cortical spreading depression

The local application of 1M potassium chloride (KCl) or tetrodotoxin (sodium channel inhibitor) induced CSD. The onset of cortical activity suppression happened after a few minutes and lasted for 15-20 minutes. To facilitate the normalization of cortical activity the cortical surface was rinsed with saline several times.

### Janelia Mouse Light Neuron Browser

We analyzed single-cell reconstruction data of the Mouse Light Neuron Browser^40^ (https://www.janelia.org/open-science/mouselight-neuronbrowser) by identifying PRF projecting M2/Cg L5 neurons, we selected neurons that have their soma in M2 or Cg area and projected to the PRF. In the browser, we could set the threshold to the axonal projection, called the in the browser “axonal endpoint”. We used 4 thresholds: >=10, 5-10, 2-5, or 1. We always precisely checked the anatomical region to uniform terminology that we used during our research (Paxinos atlas vs Neuron Browser).

### Behavior

#### Experimental protocol

After virus injection and optic fiber implantation (see above) the experimental protocol included handling (2 weeks), habituation (3 days), and the optogenetic experiment. The handling period started 1 week after the surgery. During the habituation, we placed the mouse in the experimental box and connected their optic fiber to a fiber optic patchcord (Thorlabs) and the patchcord cable (Thorlabs). The mice moved freely in the experimental box and got used to the new environment and the weight of the cable and the mouse was allowed to move passively. During the experiment phase, we photoactivated or inhibited the transfected PRF/GlyT2+ cells or their fibers in IL/Pf. The photoactivation was generated by a 473 nm DPSS laser (LRS-0473-PFM-00050-03, Laserglow Technologies, Toronto, Canada). The photoinhibition was generated by 589 nm DPSS laser (LRS-0589-GFF-00100-05, Laserglow Technologies, Toronto, Canada) The photostimulation trains were 10 s long, 5 ms light pulses at 40 Hz in PRF every 1 minute in 4 laser intensities (1, 5, 10, 15 mW). The photoinhibiton trains were 10s long, and continuous stimulation. We stimulated every condition (every optic fiber) 20 times (5-5-5-5 stimulation by intensities), and the experiment took about 30 min. During IL/Pf stimulation we used 30 Hz After the experiments, we perfused the animals and identified the location of the virus injection and the optic fibers post hoc (see Histology).

### Analysis

#### In vitro analysis

Sampling frequency was 40 kHz. Data was filtered at 3 kHz. All data were analyzed using routines developed in-house with Igor (Wavemetrics). Statistical comparisons were performed using either the Mann-Whitney test for unpaired values, or the Wilcoxon Signed-Rank test for paired sets of data, as indicated in the text. Statistical significance was set at 0.05. Results are given as mean ± s.e.m.

#### Analysis for juxtacellular recordings

All statistical analysis was performed in R Statistical Computing Software (R Core Team (2021). (R: A language and environment for statistical computing. R Foundation for Statistical Computing, Vienna, Austria. URL https://www.R-project.org/).

#### PFC photoactivation

To evaluate the effect of photoactivation the first APs following each stimulus were detected in a 40 ms window. They were distributed around ∼10 ms peak with variable spread. Only APs in the [25th percentile −1.5\*IQR, 75th percentile + 1.5\*IQR] & AP time > 0 ms range were considered evoked response. APs occurring outside of this range were considered baseline firing activity and were excluded from further analysis.

Response probabilities were calculated as the number of evoked APs/number of stimuli*100 and plotted alongside the median latency values.

#### Spontaneous frontal cortical activity changes

To determine synchronous and desynchronous periods the raw LFP signal was filtered, down-sampled, and standardized. The power spectral density of each LFP signal was calculated, and the dominant frequency was determined. To find the synchronous and desynchronous periods of the cortical LFP signal, the standard deviation of the signal amplitude was calculated in overlapping sliding windows. The size of the sliding window was equal to the length of one cycle, and the overlap was half a cycle. The threshold for synchronous and desynchronous periods was placed halfway between the median and the 25^th^ percentile.

The number of clusters was calculated in the synchronous and desynchronous periods and was statistically compared using the Wilcoxon test. Action potential clusters were determined by separating the two peaks at the minimum bimodal inter-spike interval histograms (ISI). The number of bins of the ISI was determined by the Freedman-Diaconis rule for optimal bin width; the window size was equal to the length of one cycle of the slow cortical oscillation. The mean cluster rate per LFP state was calculated by dividing the number of clusters by the length of the corresponding LFP states in every recording.

The mean firing rate (MFR) was calculated by dividing the sum number of action potentials divided by the sum lengths of the corresponding LFP states in every recording.

#### Thalamocortical neuronal activity

MFR before, during, and after stimuli were calculated as the sum number of APs divided by the sum length of each epoch. Based on the efficacy of the photo-stimulation the cells were divided into *weak* and *strong inhibition* categories. If the ratio of the number of APs +1s after and −1s before the stimulus was larger than 0.75 (number of spikes only dropped a little shortly after stimulus onset) the cell was categorized as *weak*ly inhibited, if it was smaller than 0.75 (big drop of spike numbers shortly after stimulus onset) the cell was categorized as *strong*ly inhibited.

#### Video analysis for behavior

To analyze the movements during the experiment we used Simple Mouse Tracker (SMT) downloaded from GitHub. SMT is a video tracker software based on Open CV that runs on any platform that supports Python. It is designed to track the body, head, and tail coordinates (X, Y) of a single-colour mouse over a homogeneous color background. https://github.com/joseaccruz/SimpleMouseTracker

After cleaning the tracking data (removing noise, outliers, and missing data) the before-stimulus speed and rotation of the animals were compared to the during-stimulus and -in the rotation experiments- to the after-stimulus periods.

To get the speed of the animals the Euclidean distance of the center (body) points across every 5^th^ frame was calculated and converted to mm (scaling factor: 1.6 mm/pixel). To get the rotation of the animal the signed angle of the body-head vector was calculated across every frame and the mean rotation angles across frames were plotted.

For traveled distance plots every 5^th^ frame (data point) was used (25 fps, scale: 1.6 pixel/mm). We calculated distances and average speed (mm/sec) across every 5^th^ frame. Travelled distance (mm/sec) is averaged through the last second of the “before stimulus” category and the first second of the stimulus and the last 9 seconds of the stimulus. Travelled distance “after stimulus” is not included.

For rotation, we plotted every frame (data point). To show the animal’s movement trajectory during stimuli the head-center-tail body points were plotted, separately in non-stimulated and stimulated periods. We calculated vectors (based on head-body X-Y coordination) and measured the mean angle changes of every vector. After that, we multiplied that number with fps values and divided that number by 360 to get which is the mean rotation angle during 1 sec relative to one whole circle. That value was multiplied by 100 to obtain the percentage of rotation within 1 sec in relation to a complete circle. The global angle threshold used in this file is 90. Grater rotation angles between consecutive frames are replaced by zero.

#### Statistics

In all behavioral and electrophysiological experiments, mice were randomly assigned to groups. Data points from animals with misplaced viral infections were excluded from analysis. The number of animals used in each experiment is specified in the respective figure legends. All data were checked for normality, and outliers were defined as values outside the range of Q1 - 3*IQR and Q3 + 3*IQR. All statistical analyses were performed using R Statistical Environment, Jamovi v 2.328, and Prism 8 software (GraphPad Software). For the juxtacellular analysis, the following statistical methods were applied: Mood’s median test (two-sided), Wilcoxon Signed-Rank Test (one-sided and two-sided, depending on the measure), and the Mann-Whitney test (two-sided). In vitro analysis utilized the Wilcoxon Signed-Rank Test. Behavioral data were analyzed using Friedman ANOVA with the Durbin-Conover post-hoc test, Wilcoxon Signed-Rank Test (one-sided), and the Mann-Whitney test (two-sided). Ipsilateral and contralateral axonal projection density was analyzed using a two-sided paired t-test.

## Data Availability

All relevant data supporting the findings of this study are available within the article and its supplementary materials. Additionally, source data and codes will be deposited in an appropriate public structured data repositories to ensure accessibility and future use.

Full access to the data will be granted upon publication to enable comprehensive assessment and validation of our results. In compliance with our commitment to open science, we are prepared to provide full access to the data, should it be required for the peer review.

## Acknowledgement

We thank the Light Microscopy Center, the Electron Microscopy Center and the Virus Technology Unit at the Institute of Experimental Medicine for providing support for the experiments. Authors would like to thank Krisztina Faddi for their excellent technical assistance and to Csaba Dávid for his help in quantifying the cortical input density in PRF.

## Funding

ERC Advanced Grant FRONTHAL, 742595 (LA)

European Union project within the framework of the Artificial Intelligence National Laboratory RRF-2.3.1-21-2022-00004 (LA)

Hungarian Academy of Sciences – “Lendület” Program LP2023-2/2023 (LA)

## Author contributions

Conceptualization: EB, VP, LB, LA

Methodology: EB, VP, MAD, LB, KK, LA

Investigation: EB, VP, MAD, LB, KK

Visualization: EB, VP, LB

Funding acquisition: MAD, LA

Project administration: LA

Supervision: LA

Writing – original draft: EB, MAD, VP, LB, LA

