## Supplementary material for "A cortico-subcortical loop for motor control via the pontine reticular formation": Suppl Fig 1-8

### Supplemental figures

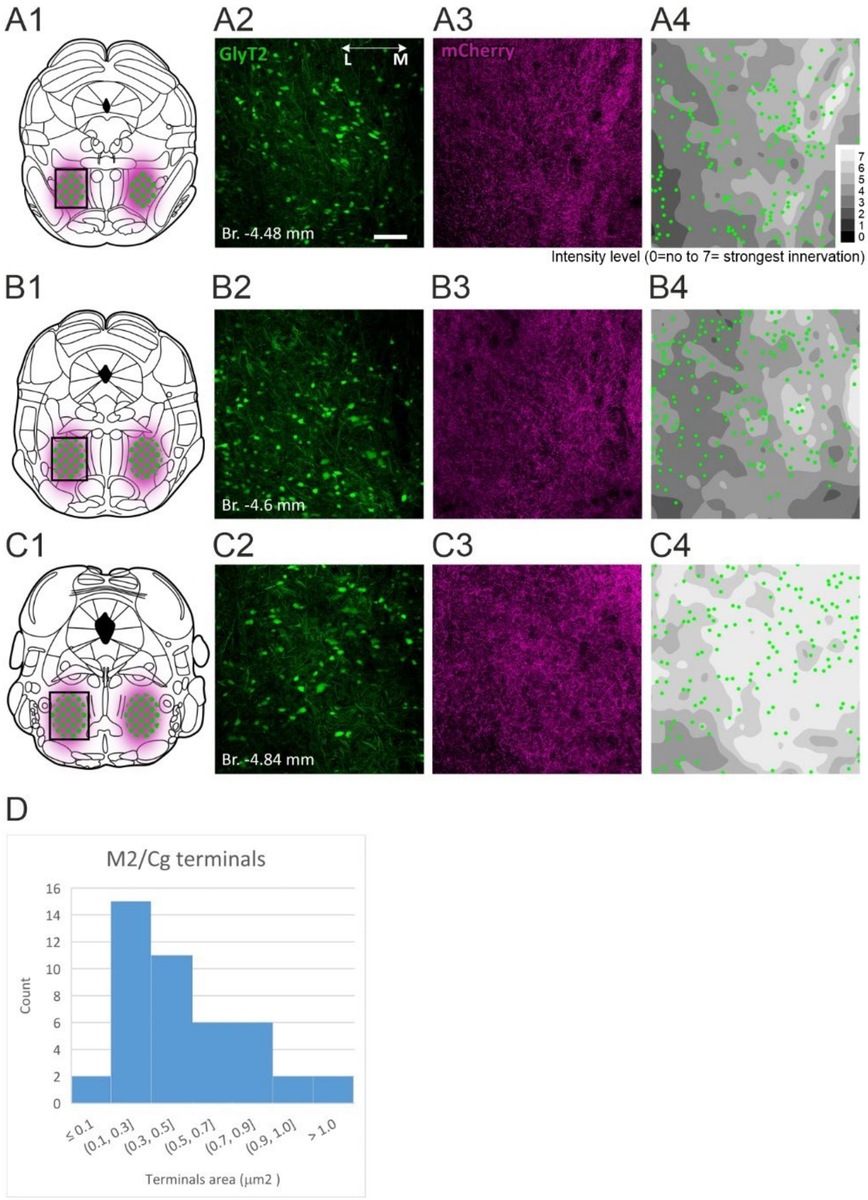

#### Figure S1. Fiber density heat maps and cortical terminal size in the PRF

A1) Schematic view of the M2/Cg L5 fibers (shady magenta area) around the PRF/GlyT2+ cells (green dots). The black rectangle indicates the position of the micrographs and heatmaps in A2-4.

A2-3) Confocal micrographs of the PRF/GlyT2+ cells (A2) and anterogradely labeled M2/Cg cortical fibers (A3) in a representative animal.

A4) Fiber density heat map (grey shading) and PRF/GlyT2+ neurons (green dots) of the same region. Higher fiber density is indicated with light grey colors.

B-C) As in A) at two more caudal levels.

D) Size Distribution of M2/Cg Terminals: the majority of M2/Cg-PRF terminals are classified as either small or medium-sized.

Scale bar: 20 μm.

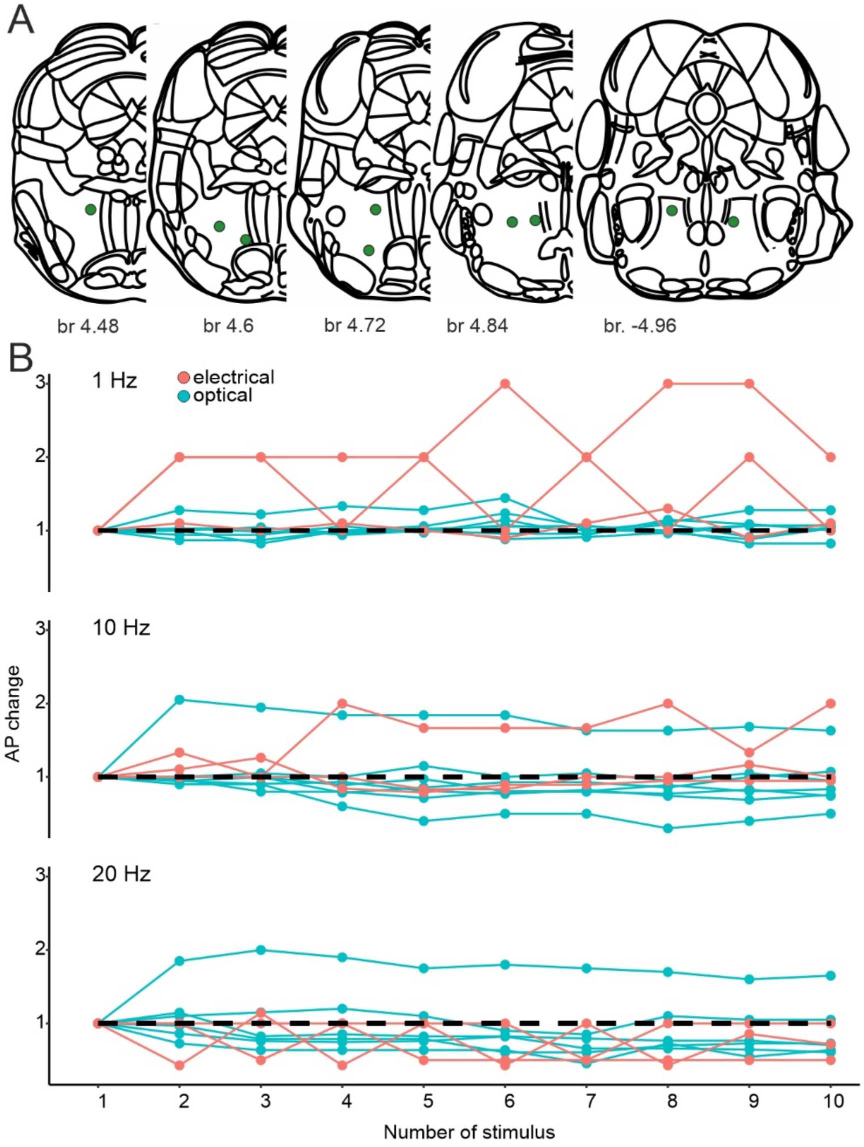


#### Figure S2. Location of the PRF/GlyT2+ neurons recorded after M2/Cg stimulation and their activity during a stimulus train

A) Post hoc identified PRF/GlyT2+ neurons (n=9 of 10 juxtacellularly recorded neuron)

B) Number of APs evoked at 1, 10, and 20 Hz during the stimulus trains in the same PRF/GlyT2+ neurons (50 ms window) normalized to the response to the first stimulus in the train (n=5 trains for each stimulus).

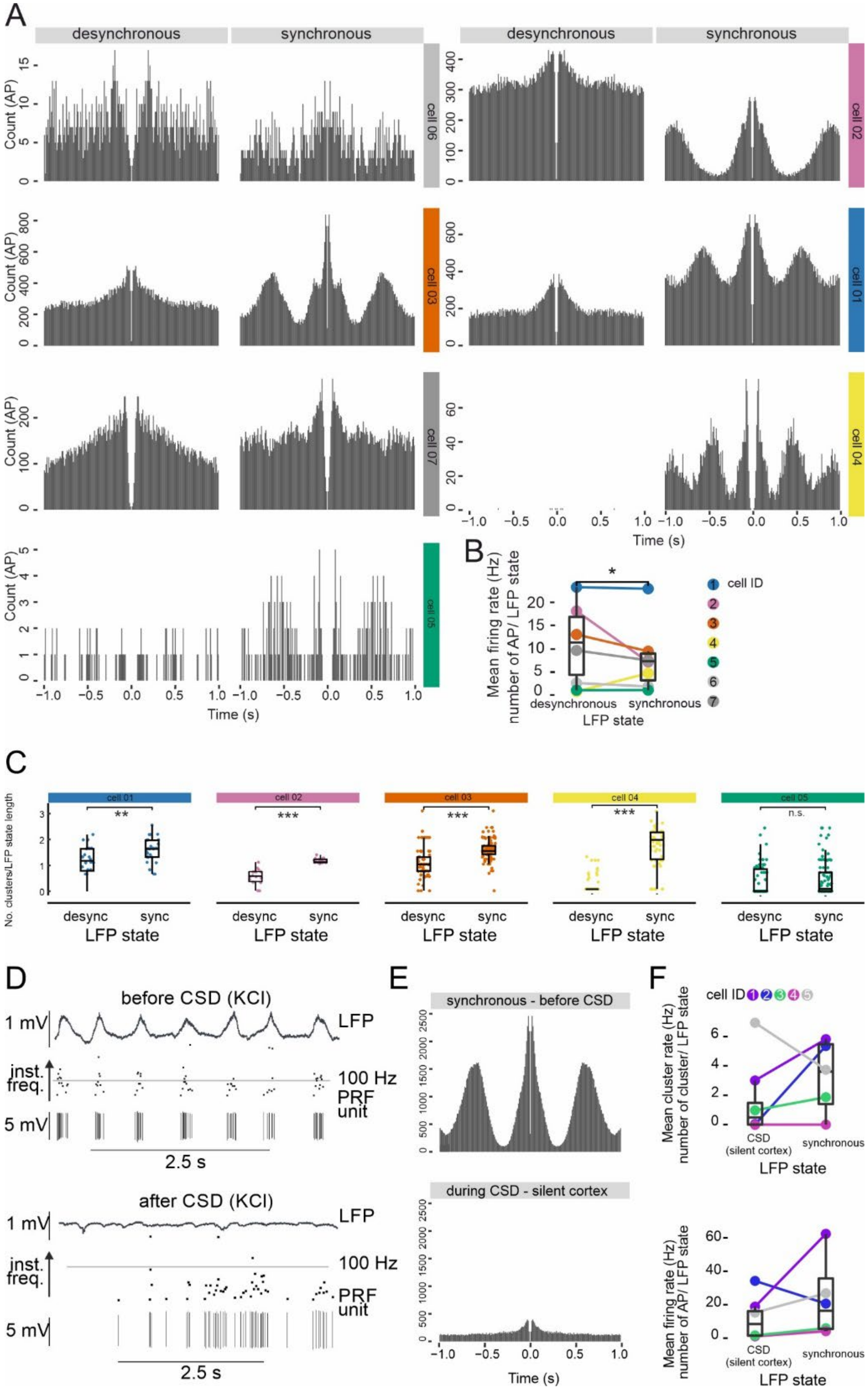
A
desynchronous
synchronous
desynchronous
synchronous
Count (AP)
cell 06
cell 02
cell 03
cell 01
cell 07
cell 04
cell 05
Time (s)
B
Mean firing rate (Hz) number of AP/ LFP state
desynchronous
synchronous
LFP state
cell ID
C
cell 01
cell 02
cell 03
cell 04
cell 05
No. clusters/LFP state length
**
***
***
***
n.s.
desync
sync
LFP state
D
before CSD (KCl)
1 mV
LFP
inst. freq.
100 Hz
PRF unit
5 mV
2.5 s
after CSD (KCl)
1 mV
LFP
inst. freq.
100 Hz
PRF unit
5 mV
2.5 s
E
synchronous - before CSD
during CSD - silent cortex
Time (s)
F
cell ID
Mean cluster rate (Hz) number of cluster/ LFP state
CSD (silent cortex)
synchronous
LFP state
Mean firing rate (Hz) number of AP/ LFP state
CSD (silent cortex)
synchronous
LFP state

#### Figure S3. Autocorrelograms, mean cluster rate, and the effect of cortical spreading depression on PRF/GlyT2+ neurons

A) Autocorrelograms during synchronized and desynchronized LFP periods (n=7)

B) Mean firing rate (number of AP/LFP state length) during synchronized and desynchronized LFP periods. Wilcoxon Signed-Rank Test, p=0.031,

C) In four out of five cells, the mean number of clusters was significantly higher during synchronous vs desynchronous LFP states. Desynchronous vs. synchronous Mann-Whitney test; cell01 p=0.0038; cell02 p<0.001; cell03 p<0.001; cell04 p<0.001; cell05 p=0.094, * 0.05<p; ** 0.01<p; *** p<0.001; n.s. - no significant difference. Source data are provided as a Source Data file.

D) Representative example of the baseline activity of a PRF neuron (top) and the changes in firing pattern following cortical inactivation using two molar KCl solution (cortical spreading depression, CSD) (bottom).

E) Autocorrelogram before CSD (top) and during CSD (bottom).

F) Mean cluster rate (number of clusters /LFP state length) during CSD (top) and mean firing rate (number of AP /LFP state length) during CSD (bottom) (n=5 neurons).

* 0.05<p; ** 0.01<p; *** p<0.001; n.s. - no significant difference. Source data are provided as a Source Data file

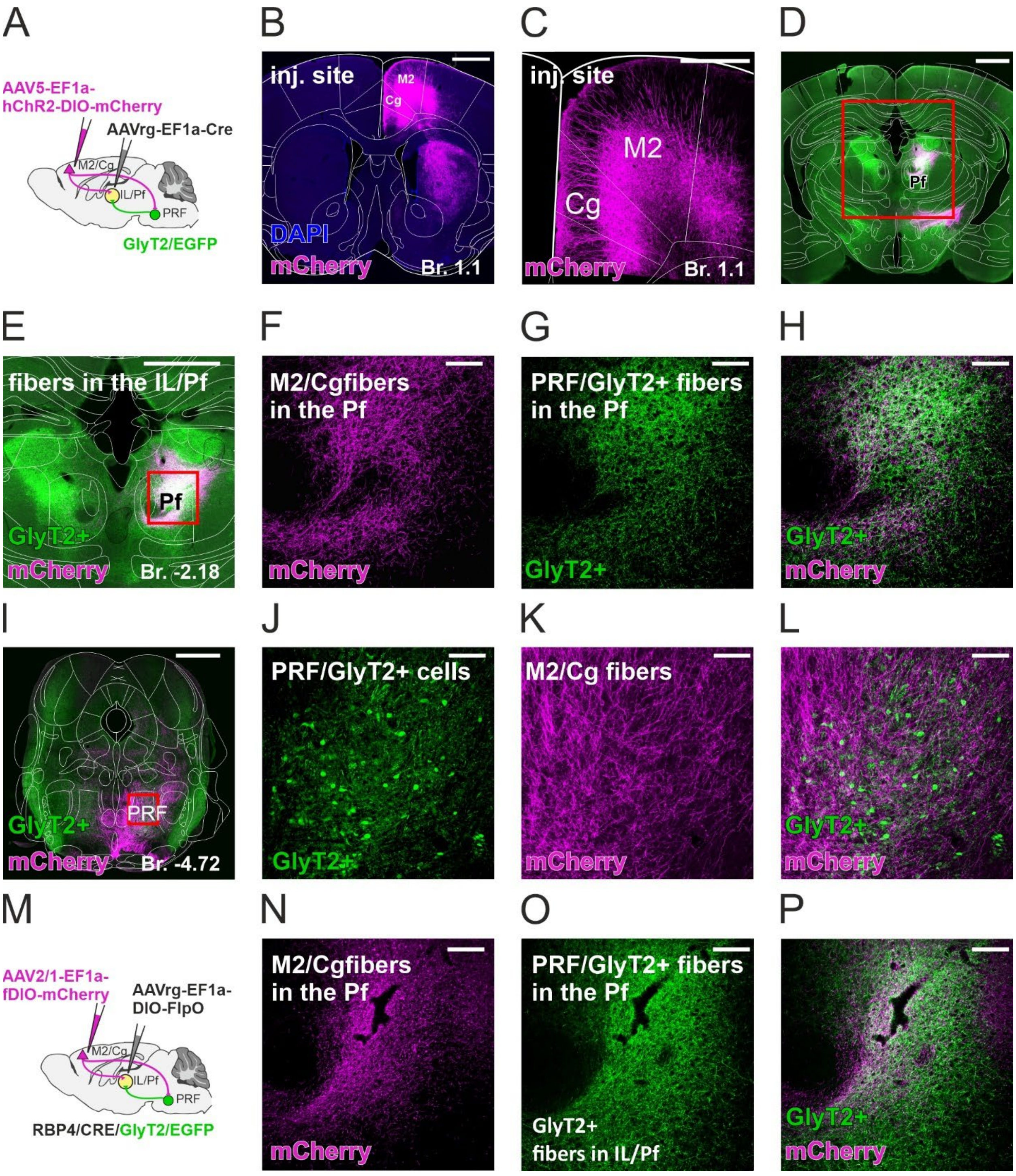
A
AAV5-EF1a-
hChR2-DIO-mCherry
AAVrg-EF1a-Cre
M2/Cg
IL/Pf
PRF
GlyT2/EGFP
B
inj. site
M2
Cg
DAPI
mCherry
Br. 1.1
C
inj. site
M2
Cg
mCherry
Br. 1.1
D
Pf
E
fibers in the IL/Pf
Pf
GlyT2+
mCherry
Br. -2.18
F
M2/Cgfibers
in the Pf
mCherry
G
PRF/GlyT2+ fibers
in the Pf
GlyT2+
H
GlyT2+
mCherry
I
PRF
GlyT2+
mCherry
Br. -4.72
J
PRF/GlyT2+ cells
GlyT2+
K
M2/Cg fibers
mCherry
L
GlyT2+
mCherry
M
AAV2/1-EF1a-
fDIO-mCherry
AAVrg-EF1a-
DIO-FlpO
M2/Cg
IL/Pf
PRF
RBP4/CRE/GlyT2/EGFP
N
M2/Cgfibers
in the Pf
mCherry
O
PRF/GlyT2+ fibers
in the Pf
GlyT2+
fibers in IL/Pf
P
GlyT2+
mCherry

#### Figure S4. Thalamus-projecting M2/Cg axon arbors in the PRF and the thalamus

A) Scheme of the second experimental design, double viral injection in GlyT2/eGFP mice

B-C) Fluorescent micrograph of the cortical injection site, low power (B) and confocal image of the cortical neurons (C).

D-E) Merged fluorescent image of the labeled thalamus-projecting M2/Cg cortical fibers (magenta) and PRF/GlyT2+ fibers (green) in the thalamus at lower (D) and higher (E) magnification. The red rectangle on D indicates the area in E and the red rectangle on E indicates the F-H position.

F-H) Confocal image of the labeled thalamus-projecting M2/Cg cortical fibers (magenta, F), of the PRF/GlyT2+ fibers (green, G) and their merged image (H) in the thalamus.

I) Composite fluorescent image of the labeled thalamus-projecting M2/Cg cortical fibers (magenta) and PRF/GlyT2+ fibers (green) in the brainstem

J-L) Confocal micrographs of the PRF/GlyT2+ cells (green, J), anterogradely labeled thalamus-projecting M2/Cg cortical fibers (magenta, K), and their merged image (L)

M) Scheme of the first experimental design (same as main Figure 5).

N-P) Confocal image of the labeled thalamus-projecting M2/Cg cortical fibers (magenta, N), of the PRF/GlyT2+ fibers (green, O) and their merged image (P) in the thalamus in the first experimental design (M).

Scale bars: B) 1 mm, C) 500 $\mu$m, D-E) 1 mm, F-H) 100 $\mu$m, I) 1 mm J-L) 100 $\mu$m, N-P) 100 $\mu$m.

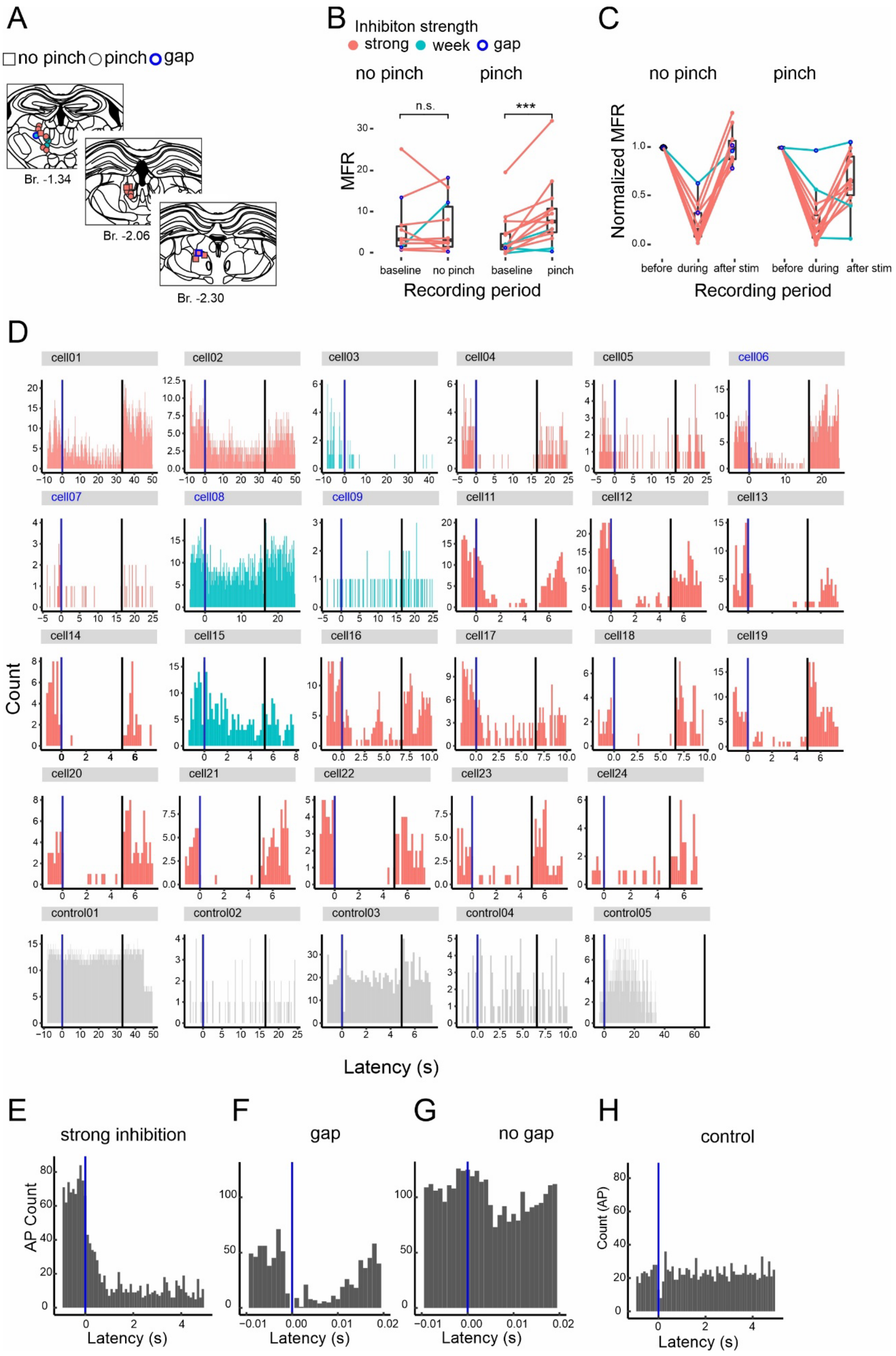

A
no pinch
pinch
gap
Br. -1.34
Br. -2.06
Br. -2.30
B
Inhibiton strength
strong
week
gap
no pinch
pinch
n.s.
***
MFR
baseline
no pinch
baseline
pinch
Recording period
C
no pinch
pinch
Normalized MFR
before
during
after stim
before
during
after stim
Recording period
D
Count
Latency (s)
E
strong inhibition
AP Count
Latency (s)
F
gap
Latency (s)
G
no gap
Latency (s)
H
control
Count (AP)
Latency (s)

#### Figure S5. Photostimulation-induced PRF/GlyT2+ mediated inhibition of IL/Pf thalamic neurons.

A) Position of the recorded IL/Pf neurons in the thalamus. Rectangles are neurons recorded with tail pinch; circles are neurons without tail pinch.

B) Comparison of mean firing rates of the IL/Pf neurons during the baseline vs before the stimulation. In the no pinch condition (n= 13 cell, left) the firing rate remains stable. In the pinch condition (right, n= 10 cell), neurons significantly increase their in response to a tail pinch. Strongly inhibited, (red circles, n=19 cell), weakly inhibited (light blue circles, n=4, see Methods) and neurons responding with a post-stimulus gap (dark blue circle, n=4) are indicated. No pinch (left) and pinch (right) conditions are shown separately. Wilcoxon Signed-Rank Test. Baseline vs. no pinch, n.s.; baseline vs. tail pinch, $p<0.001$,

C) Normalized mean firing rate (MFR) of a recorded IL thalamic neuron before, during, and after the photoactivation of PRF/GlyT2+ fibers. Labels as in B.

D) Individual PSTH of IL/Pf cell (strongly inhibited, red; weakly inhibited, blue) and and non IL/Pf thalamic control neurons (grey) before during and after the photoactivation of PRF/GlyT2+ fibers. The stimulus duration was variable (5-30 sec). Gap neurons are indicated with blue cell number fonts.

E) Population PSTH of strongly inhibited IL/Pf neuron (n=19).

F) Population PSTH of IL/Pf neuron with a post-stimulus gap in activity (n=4) in the first 20 msec.

G) Short latency population PSTH of IL/Pf neuron without a gap (n=19).

H) Population PSTH of control neurons (n=5)

* 0.05<p; ** 0.01<p; *** p<0.001; n.s. - no significant difference. Source data are provided as a Source Data file

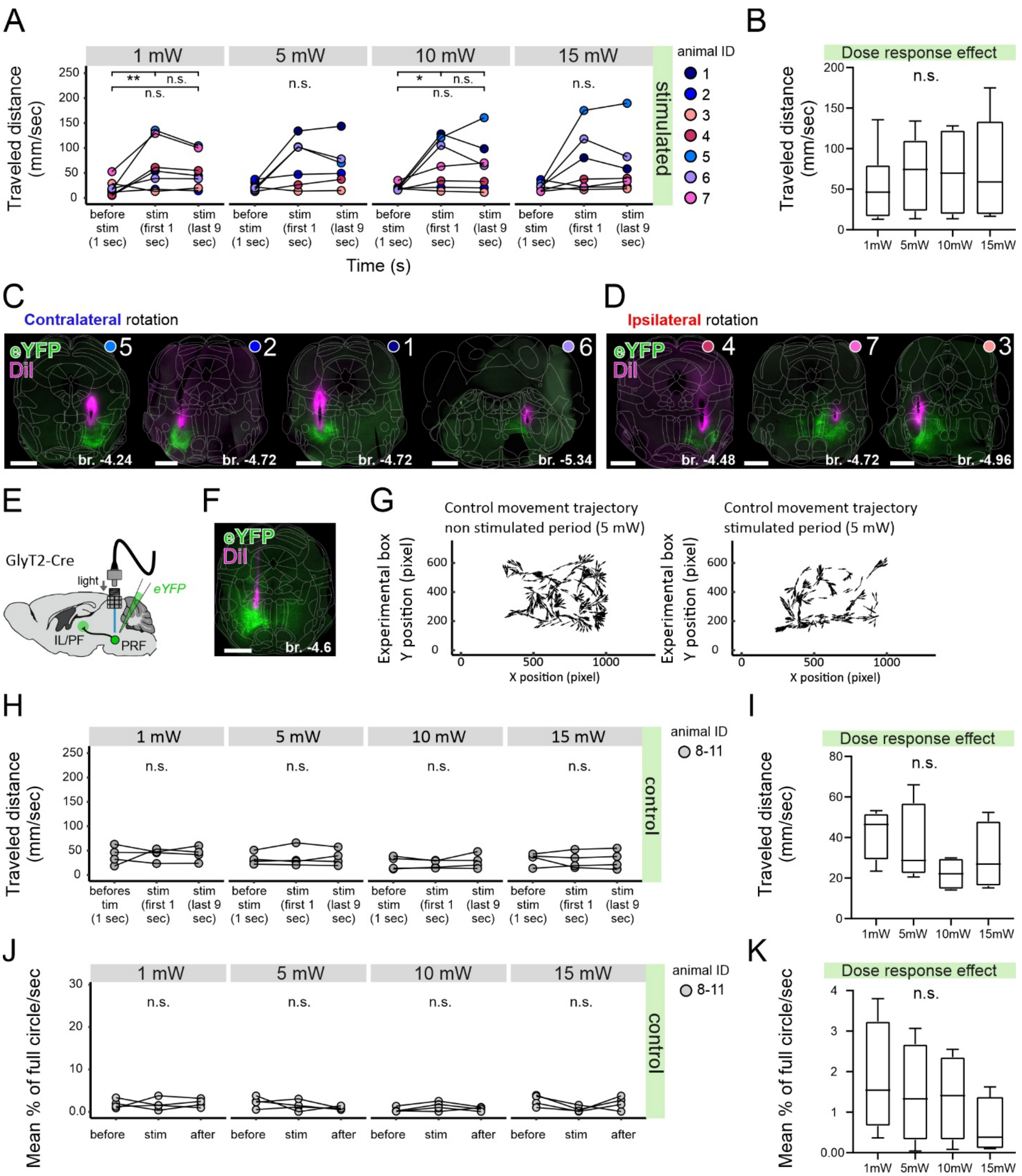

#### Figure S6. Behavioral effects of photoactivating PRF/GlyT2+ somata in experimental (ChR2) and control (EYFP) conditions

A) Comparison of the traveled distance (mm/sec) before the stimulus, in the first 1 sec and the subsequent 9 sec during the stimulus in the experimental animals (n=7). Friedman test following Durbin-Conover post-hoc, 1mW $\chi^2=7.14$, $p=0.028$, before vs first 1 sec $p=0.004$, first 1 sec vs. last 9 sec n.s., before vs. last 9 sec n.s.; 5mW $\chi^2=4.33$, n.s.; 10mW $\chi^2=9$, $p=0.050$, before vs first 1 sec $p=0.012$, first 1 sec vs. last 9 sec n.s., before vs. last 9 sec n.s.; 15mW $\chi^2=6$, n.s., numbers indicate individual animals.

B) Lack of dose-response effect at 1, 5, 10, and 15 mW laser power in experimental mice on movement initiation. Friedman test for first sec $\chi^2=4.2$, n.s.;

C) Fluorescent micrographs of the implanted optic fibers in mice responding with contralateral rotation to unilateral activation of PRF/GlyT2+ somata. Numbers/color codes are the same as for A and Figure 7F, H, I.

D) Fluorescent micrographs of the implanted optic fibers in mice responding with ipsilateral rotation to unilateral activation of PRF/GlyT2+ somata. Numbers/color codes are the same as for A and Figure 7F, H, I.

E) Experimental design of the control behavioral experiments in GlyT2-Cre mouse.

F) Fluorescent micrographs of the implanted optic fiber in representative control mice.

G) Movement trajectory in one representative control mouse during the non-stimulated period (left) and the stimulated period (right) at 5 mW.

H) Comparison of the traveled distance (mm/sec) before the stimulus, in the first 1 sec and the subsequent 9 sec during the stimulus in control animals. Friedman test 1mW $\chi^2=1.5$, n.s.; 5mW $\chi^2=0.5$, n.s.; 10mW $\chi^2=0$, n.s.; 15mW $\chi^2=0.5$, n.s.;

I) Lack of dose-response effect at 1, 5, 10, and 15 mW laser power in control mice on movement initiation. Friedman test for first sec $\chi^2=6.9$, n.s.;

J) Control animals' mean rotation angle before, during, and after stim periods. Friedman test 1mW $\chi^2=0$, n.s.; 5mW $\chi^2=1.5$, n.s; 10mW $\chi^2=1.5$, n.s.; 15mW $\chi^2=4.5$, n.s

K) Lack of dose-response effect at 1, 5, 10, and 15 mW laser power in control mice on rotation. Friedman test for first sec $\chi^2=1.2$, n.s.

* 0.05<p; ** 0.01<p; *** p<0.001; n.s. - no significant difference. Source data are provided as a Source Data file.

Scale bar: C-D) 1 mm F) 1 mm

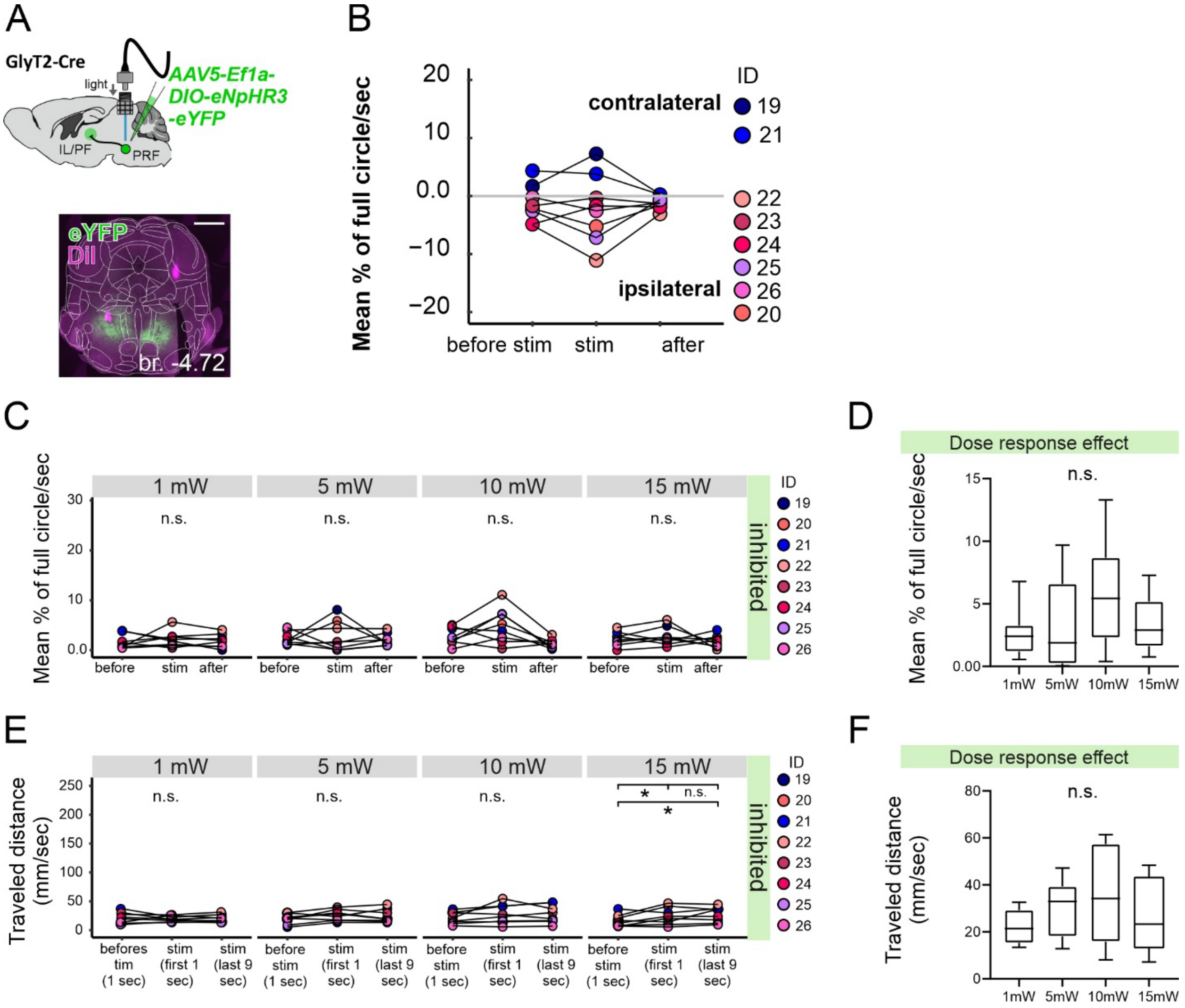


#### Figure S7. Behavioral effects of photoinhibiting PRF/GlyT2+ somata

**A)** Top: Experimental design of the photoinhibition behavioral experiments in GlyT2-Cre mouse Bottom: Fluorescent microscopic post hoc identification of implanted optic fiber and virus injection site (green) in one representative photoinhibited mouse. The appropriate optic fiber position was on the left side of the PRF labeled with DiI (magenta).

**B)** Separation of the absolute rotational values into contralateral and ipsilateral rotations. The data is displayed as the mean rotation angle before, during, and after the stimulus periods following PRF/GlyT2+ soma inhibition at all intensities.

**C)** Mean rotation angle of photoinhibited animals before, during, and after stim periods. Friedman test 1mW $\chi^2$=3, n.s.; 5mW $\chi^2$=0.25, n.s; 10mW $\chi^2$=3.25, n.s.; 15mW $\chi^2$=1, n.s

**D)** Lack of significant dose-response effect on rotation at 1, 5, 10, and 15 mW in control mice Friedman test for first sec $\chi^2$=4.2, n.s.

**E)** Comparison of the traveled distance (mm/sec) before the stimulus, in the first 1 sec and the subsequent 9 sec during the stimulus in photoinhibited animals. Friedman test following Durbin-Conover post-hoc 1mW $\chi^2$=2.25, n.s.; 5mW $\chi^2$=5.25, n.s.; 10mW $\chi^2$=1.75, n.s.; 15mW $\chi^2$=7.75, p=0.021, before vs first 1 sec p=0.039, first 1 sec vs. last 9 sec n.s., before vs. last 9 sec p=0.003

**F)** Lack of significant dose-response effect on movement initiation at 1, 5, 10, and 15 mW in control mice. Friedman test for first sec $\chi^2$=3.45, n.s.;

* 0.05<p; ** 0.01<p; *** p<0.001; n.s. - no significant difference. Source data are provided as a Source Data file.

Scale bar: A) 1 mm

s

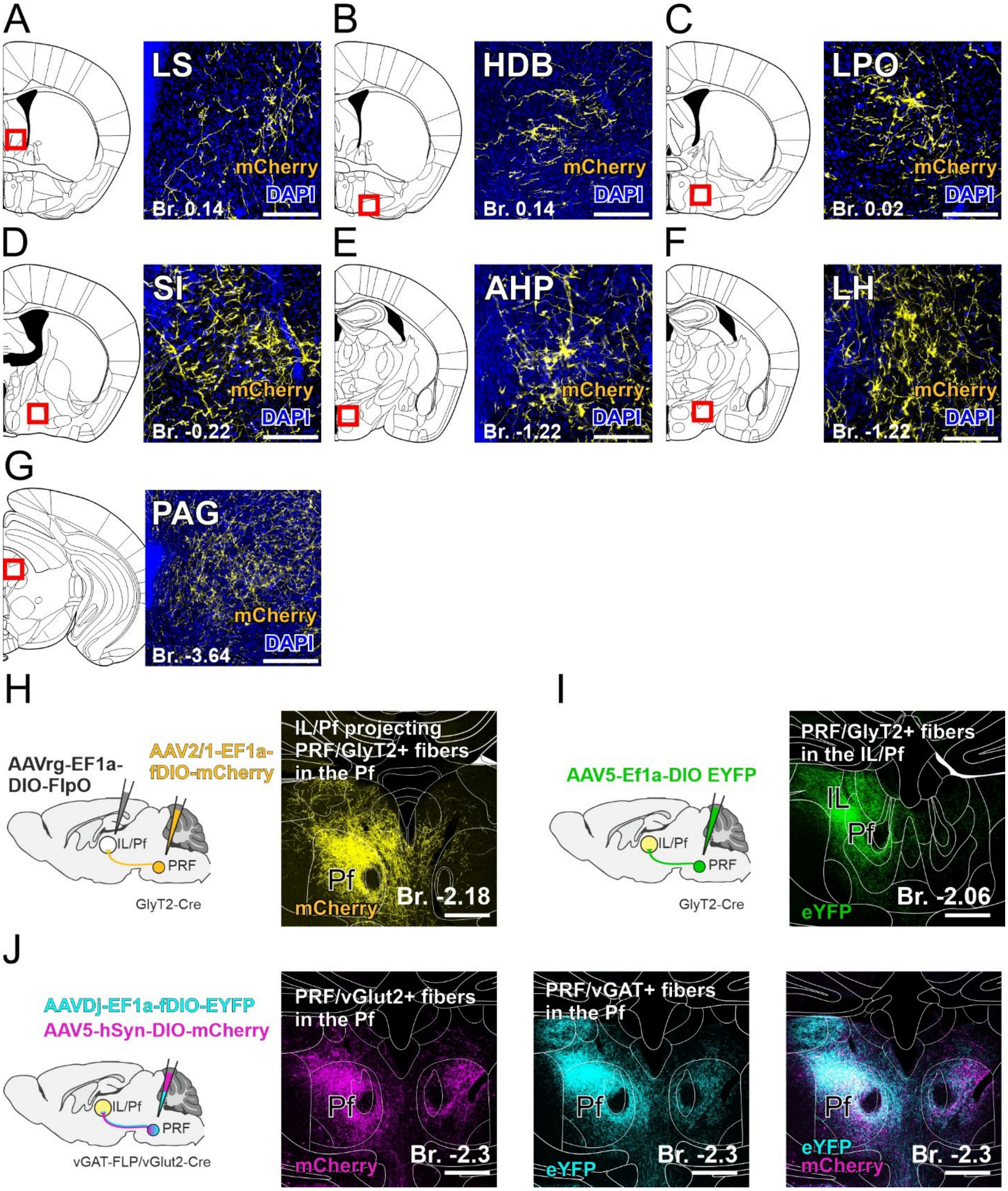
A
LS
mCherry
DAPI
Br. 0.14
B
HDB
mCherry
DAPI
Br. 0.14
C
LPO
mCherry
DAPI
Br. 0.02
D
SI
mCherry
DAPI
Br. -0.22
E
AHP
mCherry
DAPI
Br. -1.22
F
LH
mCherry
DAPI
Br. -1.22
G
PAG
mCherry
DAPI
Br. -3.64
H
AAVrg-EF1a-DIO-FlpO
AAV2/1-EF1a-fDIO-mCherry
IL/Pf
PRF
GlyT2-Cre
IL/Pf projecting PRF/GlyT2+ fibers in the Pf
Pf
mCherry
Br. -2.18
I
AAV5-Ef1a-DIO EYFP
IL/Pf
PRF
GlyT2-Cre
PRF/GlyT2+ fibers in the IL/Pf
IL
Pf
eYFP
Br. -2.06
J
AAVDj-EF1a-fDIO-EYFP
AAV5-hSyn-DIO-mCherry
IL/Pf
PRF
vGAT-FLP/vGlut2-Cre
PRF/vGlut2+ fibers in the Pf
Pf
mCherry
Br. -2.3
PRF/vGAT+ fibers in the Pf
Pf
eYFP
Br. -2.3
Pf
eYFP
mCherry
Br. -2.3

#### Figure S8. Efferents of thalamus-projecting PRF/GlyT2+.

A) Thalamus-projecting PRF/GlyT2+ axons in the lateral septum (LS).

B) Thalamus-projecting PRF/GlyT2+ axons in the nucleus of the horizontal limb of the diagonal band (HDB).

C) Thalamus-projecting PRF/GlyT2+ axons in the lateral preoptic area (LPO).

D) Thalamus-projecting PRF/GlyT2+ axons in the subtantia innominate (SI).

E) Thalamus-projecting PRF/GlyT2+ axons in the anterior hypothalamus (AHP).

F) Thalamus-projecting PRF/GlyT2+ axons in the lateral hypothalamus (LH).

G) Thalamus-projecting PRF/GlyT2+ axons in the periaqueductal grey (PAG).

H) Scheme of double conditioned viral tracing for investigating the thalamus-projecting PRF/GlyT2+ cells' efferents in GlyT2-Cre mice (right) and their axons in the thalamus (left).

I) Experimental design to label all PRF/GlyT2+ cells (left) and their axons in the thalamus (right).

J) Scheme of mixed viral injection into PRF in vGAT-Flp/vGlut2-Cre (right) and their vGlut2+ fibers (magenta); PRF/vGAT+ fibers (cyan) and merged confocal image of the vGlut2+ and vGAT+ fibers (left) in the Pf.

Scale bars: A-G) 200 μm, H-J) 500 μm.

### Supplemental tables

| Co-innervation of PRF and IL/Pf by single L5 neurons | | | | | |
|---|---|---|---|---|---|
| **DOI** | **Soma's position** | **Entire name** | **PRF axonal endpoint** | **IL/Pf axonal endpoint** | **striatal axonal endpoint** |
| AA0179 | MOs5 | Secondary motor area layer 5 | 1 | 2 | + |
| AA0845 | ACAv5 | Anterior cingulate area ventral part layer 5 | 1 | 0 | - |
| AA0115 | MOs5 | Secondary motor area layer 5 | 2 | 4 | + |
| AA0764 | ACAd5 | Anterior cingulate area dorsal part layer 5 | 2 | 3 | + |
| AA0796 | ACAv5 | Anterior cingulate area ventral part layer 5 | 3 | 0 | - |
| AA0882 | MOs5 | Secondary motor area layer 5 | 3 | 0 | - |
| AA1544 | MOs5 | Secondary motor area layer 5 | 3 | 4 | - |
| AA0181 | MOs5 | Secondary motor area layer 5 | 4 | 44 | + |
| AA0415 | MOs5 | Secondary motor area layer 5 | 4 | 2 | + |
| AA0576 | MOs5 | Secondary motor area layer 5 | 4 | 1 | + |
| AA0792 | MOs5 | Secondary motor area layer 5 | 4 | 20 | + |
| AA0182 | MOs5 | Secondary motor area layer 5 | 5 | 14 | + |
| AA0250 | MOs5 | Secondary motor area layer 5 | 6 | 1 | + |
| AA0780 | MOs5 | Secondary motor area layer 5 | 6 | 7 | + |
| AA1538 | MOs5 | Secondary motor area layer 5 | 6 | 0 | - |
| AA0114 | MOs5 | Secondary motor area layer 5 | 7 | 5 | + |

| | | | | | |
|---|---|---|---|---|---|
| AA0791 | MOs5 | Secondary motor area layer 5 | 8 | 6 | + |
| AA0180 | MOs5 | Secondary motor area layer 5 | 9 | 0 | - |
| AA1541 | MOs5 | Secondary motor area layer 5 | 12 | 0 | - |
| AA0772 | MOs5 | Secondary motor area layer 5 | 19 | 31 | + |
| AA0788 | MOs5 | Secondary motor area layer 5 | 19 | 6 | + |
| AA0261 | MOs5 | Secondary motor area layer 5 | 37 | 36 | + |
| AA0245 | MOs5 | Secondary motor area layer 5 | 39 | 34 | + |

##### Table S1. Co-innervation of PRF,IL/Pf and the striatum by single L5 neurons

Neuron database based on Janelia Mouse Browser. DOI: Janelia ID for neurons. Soma's position name was determined by Janelia's nomenclature PRF and IL/Pf axonal endpoint means how many afferents the PRF has from M2/Cg. In the striatal axonal endpoint column, the +/- indicates whether there was M2/Cg's afferent.

#### Legend for supplemental videos

**Video S1 –** Contralateral turning during the PRF/GlyT2+ fiber stimulation in the Pf at 10 mW including the before and after stimulus period. The light on the upper right side indicates the stimulation period.

**Video S2 –** Contralateral turning during the PRF/GlyT2+ cell stimulation at 5 mW including the before and after stimulus period. The light on the right side indicates the stimulation period.

**Video S3 –** Ipsilateral turning during the PRF/GlyT2+ cell stimulation at 10 mW including the before and after stimulus period. The light on the right side indicates the stimulation period.

**Video S4 –** Control animal during the PRF/GlyT2+ cell stimulation at 10 mW including the before and after stimulus period. The light on the right side indicates the stimulation period.

Video S5 - Contralateral turning during the PRF/GlyT2+ soma photoinhibiton at 5 mW including the before and after stimulus period. The light on the upper side indicates the stimulation period.
